# Regulatory scope shapes the adaptive landscapes of bacterial transcription factor binding sites

**DOI:** 10.64898/2026.08.11.744152

**Authors:** Cauã Antunes Westmann, Andreas Wagner

## Abstract

Transcription factors (TFs) span a regulatory hierarchy from local regulators that control one or few genes to global regulators that regulate hundreds. Local TFs typically operate through few, highly specific binding sites; global TFs through many of varying affinity. Whether this difference in TF biology systematically affects TFBS evolution is unclear. Here, we address this question by studying experimentally mapped adaptive landscapes of transcription factor binding sites (TFBSs) for five bacterial TFs that differ in their regulatory scope — TetR (local), LasR (semi-global), and CRP, Fis, and IHF (global). All landscapes are rugged and epistatic, but three topographic properties vary systematically with regulatory scope. First, mean mutational robustness increases from local to global TFs. Second, high-regulation-strength peaks are clustered in the local landscape but dispersed in the global ones. Third, adaptive evolution reaches high adaptive peaks more readily in local landscapes. All five landscapes also harbor many evolvability-enhancing (EE) mutations, which increase the likelihood that subsequent mutations are beneficial. The fraction of these mutations is approximately an order of magnitude higher than previously reported for a protein landscape. Populations experiencing EE mutations consistently reach higher regulation strength, an effect that is strongest in the local TetR landscape. Together, these results identify regulatory scope as an organising axis of TFBS adaptive landscapes.

## Introduction

Gene regulation helps organisms respond to environmental signals, and its evolution plays a central role in generating phenotypic diversity ^1,2^. Gene regulation requires transcription factors (TFs), proteins that regulate gene expression by binding to specific DNA sequences — transcription factor binding sites (TFBSs) – thereby activating or repressing a gene’s transcription. TFs form hierarchical gene regulatory networks (GRNs) ^3^ in which local regulators near the bottom of the hierarchy control only one or a few genes, whereas a few global regulators at the top control the expression of many target genes — including genes that themselves encode transcription factors.

Because TFs often regulate multiple genes, mutations in TFs can have widespread pleiotropic effects. Regulatory evolution therefore frequently proceeds through mutations in TFBSs, which can modulate the expression of one gene with no deleterious consequences for other genes ^2,4^. Such regulatory evolution has contributed to diverse adaptive traits across taxa. For example, in snakes evolutionary loss of limbs has been linked to deletions in limb-specific regulatory DNA of the Shh gene^5^. In butterflies, wing-pattern diversification involves mutations in TFBSs regulating the genes *optix* and *WntA* ^6^. In *Escherichia coli*, regulatory mutations upstream of the *csgD* gene alter biofilm formation ^7^.

Transcription factors are commonly classified by their *regulatory scope*, the size of their regulon — the number of genes whose expression they influence^8–10^. By this definition, local transcription factors regulate one or a few genes, semi-global transcription factors regulate intermediate numbers of genes, and global transcription factors regulate many genes and pathways. Regulon size does not necessarily reflect the number of genomic sites a transcription factor directly binds, because regulatory hierarchies can amplify the number of genes that a factor regulates directly or indirectly ^11^. The number of directly bound DNA sites provides a complementary view of regulatory scope that is especially relevant to transcriptional promiscuity: because promiscuous factors recognise many sequences, new binding sites — and thus new regulatory connections — could arise more readily during evolution for such TFs^12–14^.

An example of a local regulator is TetR. It represses a single tetracycline-resistance operon from a single operator region^15^. By contrast, the quorum-sensing regulator LasR of *Pseudomonas aeruginosa* is semiglobal. It affects the expression of more than 300 genes, corresponding to roughly 5% of the *P. aeruginosa* genome^16,17^, even though it binds only approximately 35 promoter regions that control up to 74 genes ^18,19^. Global regulators include the *E.coli* cAMP receptor protein (CRP) which binds at least 254 genomic sites^20,21^, the factor for inversion stimulation (Fis), which binds approximately 900 sites ^22–24^, and the integration host factor (IHF), which binds near approximately 30% of predicted operons ^25,26^. Whether these differences in regulatory scope translate into systematic differences in TFBS evolution is unknown.

An adaptive landscape of TFBSs maps binding-site genotypes to regulatory phenotypes related to gene expression level ^27–30^. In the landscapes we study here, this phenotype is regulation strength, defined as the ability of a TFBS to modulate the expression of a reporter gene’s expression by binding its cognate TF. Each TFBS occupies a location in the sequence space underlying such a landscape, and its elevation corresponds to experimentally measured regulation strength. We use regulation strength as a proxy for fitness, which reflects the assumption that strong regulation can be beneficial. We use regulation strength as a proxy for fitness, which reflects the assumption that strong regulation can be beneficial. This has been demonstrated for individual genes across organisms as diverse as bacteria, yeast, plants, and humans^31–34^. Relatedly, the binding affinity of a TF to its TFBS is subject to natural selection in multiple species ^31,35–37^. More broadly, regulation strength is subject to selection whenever insufficient or excessive expression imposes fitness costs. Because the relationship between regulation strength and fitness can be non-monotonic, stronger regulation is not universally beneficial in nature. However, our assumption provides a well-defined selective regime that allows us to compare the topography of different TFBS landscapes.

This topography can also constrains how populations evolve in sequence space ^27,38,39^. Peaks are TFBS genotypes whose regulation strength exceeds that of all their single-mutant neighbours — DNA sequences that differ from the focal genotype by one nucleotide. The highest such genotype is the global peak; all others are local peaks. Under directional selection for increased regulation strength, a population can ascend a high fitness regions and reach a peak. In a smooth landscape, this peak is also the *global peak* with the highest regulation strength among all genotypes. In rugged landscapes with many peaks, a population may become trapped at suboptimal *local peaks* because all one-step mutations away from such peaks reduce regulation strength and are therefore selected against. Landscape ruggedness — the number and location of peaks — can therefore affect how readily evolving populations can reach a global peak ^40,41^.

Here we model adaptive evolution through adaptive walks, in which an evolving population is monomorphic most of the time, that is, it consists of a single genotype, and periodically experiences mutations that rapidly sweep through the population, and that effectively constitute a single step through an adaptive landscape. This modeling framework is appropriate whenever beneficial mutations are rare, such as in the evolution of TFBSs, which are short and thus represent small mutational targets ^41–44^. In this framework, it is also natural to define the evolutionary *accessibility* of a peak as the probability that an adaptive walk starting from a given genotype reaches the peak. Accessibility depends on the existence of mutational paths through a genotype space in which each mutation increases regulation strength^40,41^.

One important property of TFBS genotypes is mutational robustness: the capacity of a sequence to maintain its regulatory ability despite mutations ^45–47^. Robustness is fundamentally a property of individual genotypes, but its distribution across a landscape can also help to characterize how tolerant a TFs gene regulation ability is to mutations in general. Robustness is closely linked to evolvability, the ability to generate adaptive variation through mutation ^46^, because buffering mutational effects can both preserve an existing phenotype and enhance access to novel phenotypes ^46,47^. Although prior work has examined this robustness–evolvability relationship in individual TFBS landscapes ^48^, it remains unknown whether this relationship varies systematically with the regulatory scope of TFs.

A complementary dimension of evolutionary potential is captured by evolvability-enhancing (EE) mutations ^49^. An EE mutation creates a genetic background in which subsequent mutations are more likely to be adaptive than before the EE mutation. Such a mutation may spread through direct selection on its own fitness effects, and in doing so facilitate adaptive evolution^49–51^. EE mutations have been described in RNA and protein adaptive landscapes ^49–51^. It is unknown whether they also occur in TFBS landscapes, and whether their prevalence depends on regulatory scope.

Here, we analyse the in vivo adaptive landscapes of TFBSs for five bacterial TFs spanning the local-to-global regulatory hierarchy: TetR ^30^ (local), LasR ^18^ (semi-global), and CRP, Fis, and IHF ^29^ (global). Using Sort-Seq ^52^, a high-throughput method that links a TFBS genotype to its ability to regulate gene expression in vivo, we measured the regulation strength of thousands of binding-site variants per TF and compared landscape topography, peak accessibility, mutational robustness, and EE-mutation prevalence across TFs. We find that key features of TFBS adaptive landscapes vary systematically with regulatory scope: mean mutational robustness increases from local to global TFBS landscapes, high-regulation-strength peaks are more clustered in local landscapes than in global ones, and adaptive walks more readily reach high peaks in local landscapes. EE mutations are abundant in all landscapes. They facilitate adaptive evolution by increasing the regulation strength that an evolving population can reach, and do so most strongly in the local TetR landscape. Together, these results identify regulatory scope as an organising axis of TFBS adaptive landscapes, and link gene regulatory network architecture to the evolutionary potential of cis-regulatory sequences.

## Results

### Mapping adaptive landscapes of TFBSs

The Sort-Seq method links TFBS genotype to gene-expression phenotype by coupling libraries of TFBS variants to a fluorescent reporter gene, sorting individual cells into multiple fluorescence bins through fluorescence-activated cell sorting (FACS), and quantifying variant frequencies across fluorescence bins by deep sequencing ^29,30,53^ (see **Methods** for details). For our Sort-Seq assay, we placed each TFBS variant upstream of a green fluorescent protein (GFP) reporter gene regulated by its cognate TF. Variants that confer weak regulation produce high GFP expression and are enriched in high-fluorescence bins, whereas variants that confer strong regulation produce low GFP expression and are enriched in low-fluorescence bins. Deep sequencing of each bin therefore makes it possible to estimate the expression level, and hence the regulation strength, of each TFBS variant.

Using this method, we mapped TFBS adaptive landscapes for all five TFs (**Fig. 1**)^29,30^.For each TF, we randomised the eight most informative positions within its binding motif, yielding a maximal diversity of 4⁸ = 65,536 variants. We then inferred the expression level of each variant from its distribution across fluorescence bins and defined its regulation strength as the inverse of GFP expression, normalised from 0 (weakest regulation) to 1 (strongest regulation) (**Supplementary Methods Sections 3, 4**).

**Figure 1.**
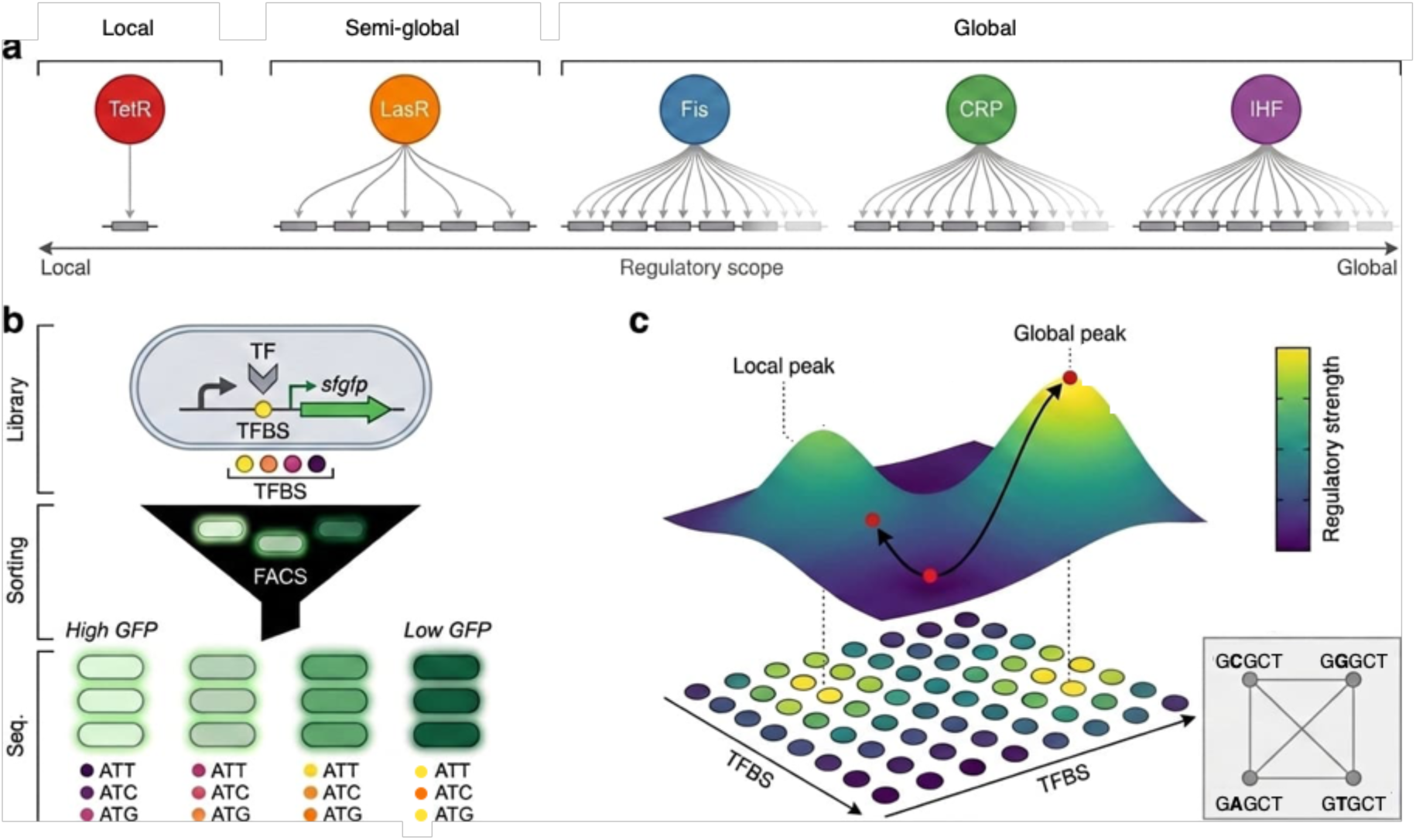
Conceptual overview of TFBS adaptive landscapes across the local-to-global regulatory hierarchy. a. Bacterial transcription factors analysed in this study: TetR (local), LasR (semi-global), and CRP, Fis, and IHF (global). Increasing numbers of regulated operons (grey boxes) illustrate increasing regulatory scope. The same color code is used for each TF throughout the manuscript. b. Sort-Seq mapping of TFBS adaptive landscapes. We cloned libraries of TFBS variants upstream of an *sfgfp* reporter and sorted them into fluorescence bins by FACS according to GFP expression. We used deep sequencing of each bin to infer a normalised regulation-strength score for each variant, scaled from 0 (weakest regulation) to 1 (strongest regulation). (see Methods) Light-to-dark green shading indicates increasing regulation strength. c. Schematic representation of a TFBS adaptive landscape. Genotypes are connected by single-nucleotide mutations (inset), and landscape elevation corresponds to regulation strength. Surface colour indicates regulation strength, from low (purple) to high (yellow). Peaks represent local maxima in regulation strength. High peaks exceed the regulation strength of the wild-type TFBS. The illustrated adaptive walks (black arrows; red circles) show trajectories that climb the landscape and can terminate either at a local peak or at the global peak. We restricted analyses to the largest connected component of each genotype network, yielding landscapes containing 17,765 (TetR), 43,087 (LasR), 31,975 (CRP), 43,222 (Fis), and 41,325 (IHF) genotypes.

After quality filtering (**Supplementary Methods Section 4**), we retained 17,765 binding site genotypes for TetR, 43,087 for LasR, 31,975 for CRP, 43,222 for Fis, and 41,325 for IHF. We represented an adaptive landscape as a network in which nodes correspond to TFBS variants, and edges connect variants that differ by a single nucleotide. We restricted analyses to the largest connected component^54^ of each network (landscape), which comprises approximately 97% of genotypes in each landscape. The TetR, CRP, Fis, and IHF data derive from two previous studies ^29,30^, whereas we generated the LasR dataset specifically for this study (**Supplementary Methods Section 2**).

Each landscape spanned a broad range of regulation strengths (**Supplementary Fig. S1**). For the semi-global TF LasR and the global TFs CRP, Fis, and IHF, this experimentally measured range closely matched the distribution of regulation strengths predicted *in silico* for curated genomic binding sites^55^ (**Supplementary Fig. S2**). This agreement suggests that our experimental assay and TFBS libraries capture regulation-strength ranges relevant to natural TFBS evolution. An equivalent comparison is not possible for TetR because only two cognate genomic TFBSs are known^15^.

### Mutational effects and robustness vary with regulatory scope

We first quantified the distribution of single-nucleotide mutational effects on regulation strength across the five landscapes. This distribution differs systematically among TF classes (**Supplementary Fig. S4**): It is broadest (with highest variance) for the local regulator TetR, narrowest (with lowest variance) for the global regulators (CRP, Fis, IHF), and intermediate for the semi-global LasR. This pattern persists within each regulation-strength class: mutations in TetR produce larger changes in expression than those in CRP, Fis, or IHF at low, medium, and high regulation strengths alike (**Supplementary Fig. S4**). Pairwise two-sample Kolmogorov–Smirnov (KS) tests reject the null hypothesis that mutational effects in each pair of landscapes are drawn from the same distribution (all P < 10⁻⁵⁶; full sample sizes and pairwise statistics in **Supplementary Fig. S4**), with KS statistics ranging from D = 0.140 (TetR vs Fis) to D = 0.384 (CRP vs LasR).

We next quantified mutational robustness for each genotype *g* as 1 − *R*_avg_(*g*), where *R*_avg_(*g*) is the mean absolute change in regulation strength between *g* and its single-nucleotide neighbours. Higher values of 1 − *R*_avg_(*g*) indicate greater robustness (**Fig. 2a**). Across all landscapes, most mutations reduce regulation strength, just like most mutations reduce fitness in multiple biological systems^56^, including TFBSs^29,30,45,57^. Within each landscape, robustness shows a non-monotonic relationship with regulation strength — genotypes of intermediate strength are the most robust, whereas low-and high-strength genotypes are more sensitive to mutation (**Fig. 2c**). This observation is consistent with previous observations using a different robustness metric on σ⁷⁰ promoters^58^.

**Figure 2.**
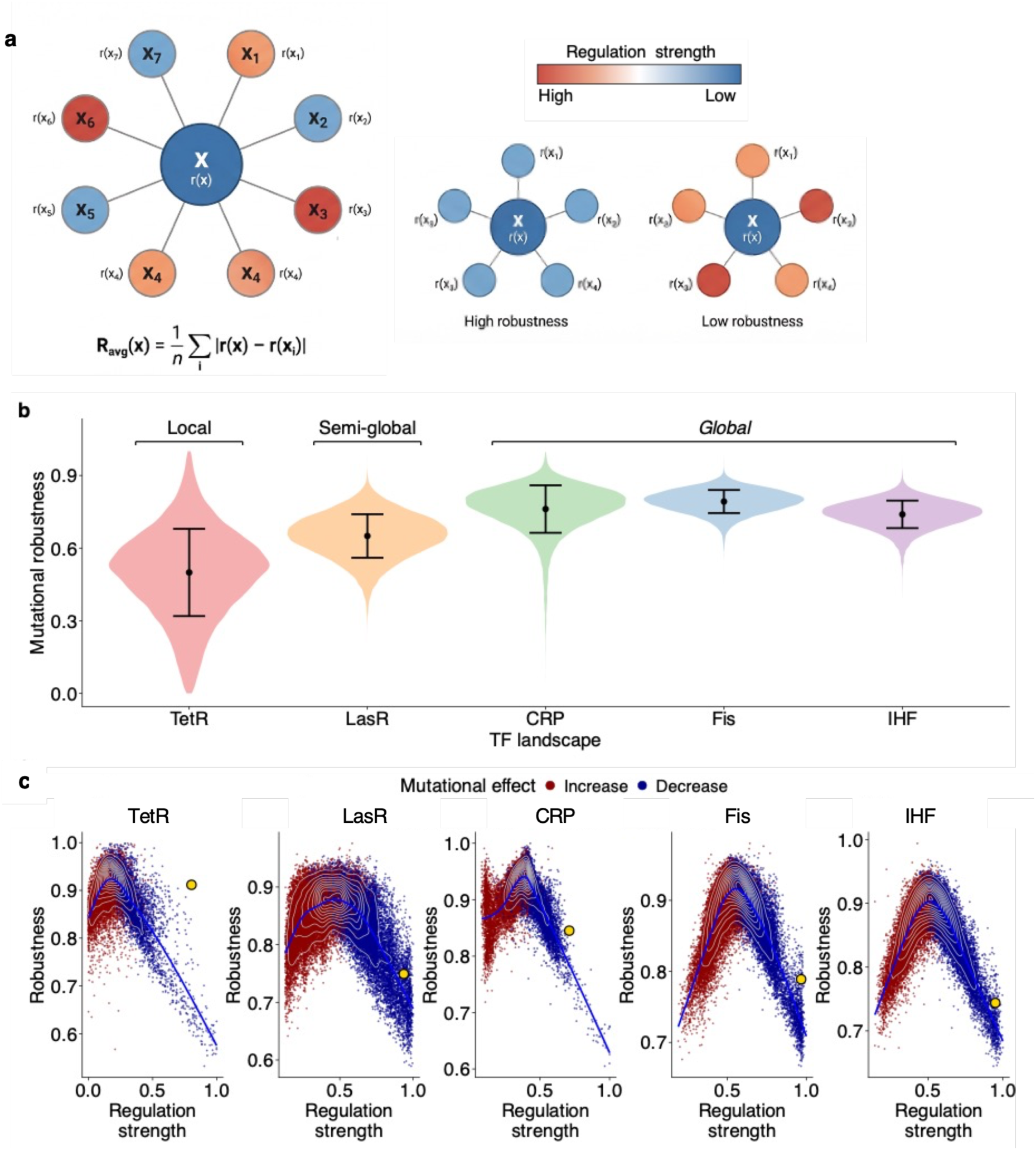
Mutational effects and robustness vary systematically with regulatory scope. a Illustration of the robustness metric. For a focal genotype, mutational sensitivity is quantified as the mean absolute change in regulation strength between the focal genotype and its single-nucleotide neighbours, *R*_avg_(*g*) (**Methods**). Mutational robustness is reported as 1 − *R*_avg_(*g*), so higher values indicate greater robustness. **b Distribution of mutational robustness across the five landscapes.** Brackets above the violin plots group landscapes by regulatory scope (Local: TetR; Semi-global: LasR; Global: CRP, Fis, IHF). Mean robustness values are 0.485 (TetR), 0.651 (LasR), 0.763 (CRP), 0.792 (Fis), and 0.739 (IHF). Differences among landscapes are significant under a Kruskal–Wallis test rejecting the null hypothesis that robustness values are drawn from the same distribution across landscapes (*H* = 80,129.4, df = 4, P < 2.2 × 10⁻¹⁶; Dunn’s post-hoc test with Bonferroni correction, all pairwise P < 0.001). Sample sizes are n = 17,765 (TetR), 43,087 (LasR), 31,975 (CRP), 43,222 (Fis), 41,325 (IHF) genotypes. **c Robustness as a function of regulation strength for each TF.** Each point represents one genotype and is coloured by the sign of the mean effect of mutations on regulation strength, *ΔS_R_*: blue indicates that mutations increase regulation strength on average, whereas red indicates that mutations decrease regulation strength on average. White contour lines show two-dimensional kernel density estimates. The blue curve is an estimated regression fit to the data. Yellow circles mark the wild-type genotype in each landscape, with the following wild-type regulation strengths: TetR = 0.804, LasR = 0.936, CRP = 0.710, Fis = 0.967, and IHF = 0.949. Across all five landscapes, robustness is highest at intermediate regulation strength and lower at both extremes. Sample sizes are as in panel b. The distribution of single-nucleotide mutational effects on regulation strength is shown in **Supplementary Fig. S4**.

Peak genotypes are less robust than non-peak genotypes in every landscape (two-sided Wilcoxon rank-sum test, P < 2.2 × 10⁻¹⁶ in each; N_peaks = 2,068–2,450, N_other = 13,388– 40,894). Because such large samples render even negligible differences significant, we quantified the size of this difference with Cliff’s delta^59^ (δ; **Methods**), which ranged from −0.45 in TetR to −0.88 in LasR (**Supplementary Fig. S5**, **Supplementary Methods 12.2**). The negative sign indicates lower robustness for peaks than for non-peaks, and the magnitudes correspond to moderate (TetR) to large (LasR) effects

Beyond these within-landscape patterns, landscape-mean robustness also varies systematically with regulatory scope (**Fig. 2b**). Mean robustness is lowest for TetR (0.485), intermediate for LasR (0.651), and highest for the global regulator landscapes (IHF 0.739, CRP 0.763, Fis 0.792; Kruskal–Wallis^60^ test rejecting the null hypothesis that robustness values are drawn from the same distribution across landscapes: *H* = 80,129.4, df = 4, *P* < 2.2 × 10⁻¹⁶; Dunn’s post-hoc tests^61^ with Bonferroni correction, all pairwise P < 0.001).

### High peaks are more clustered in local but dispersed in global landscapes

All five landscapes are rugged (**Table 1; Supplementary Methods Section 7**), but the spatial organisation of high peaks differs systematically across TF classes (**Fig. 3**). Local peaks comprise 12.2% of genotypes in TetR — more than double the 5.4–6.7% in LasR, CRP, Fis, and IHF. High peaks (those exceeding wild-type regulation strength; **Supplementary Methods Section 7.2**) are rare in all landscapes (0.2–0.5% of genotypes). For comparison, peak fractions in this range place all five landscapes among the most rugged empirical adaptive landscapes characterised to date^27,62,63^.

**Figure 3.**
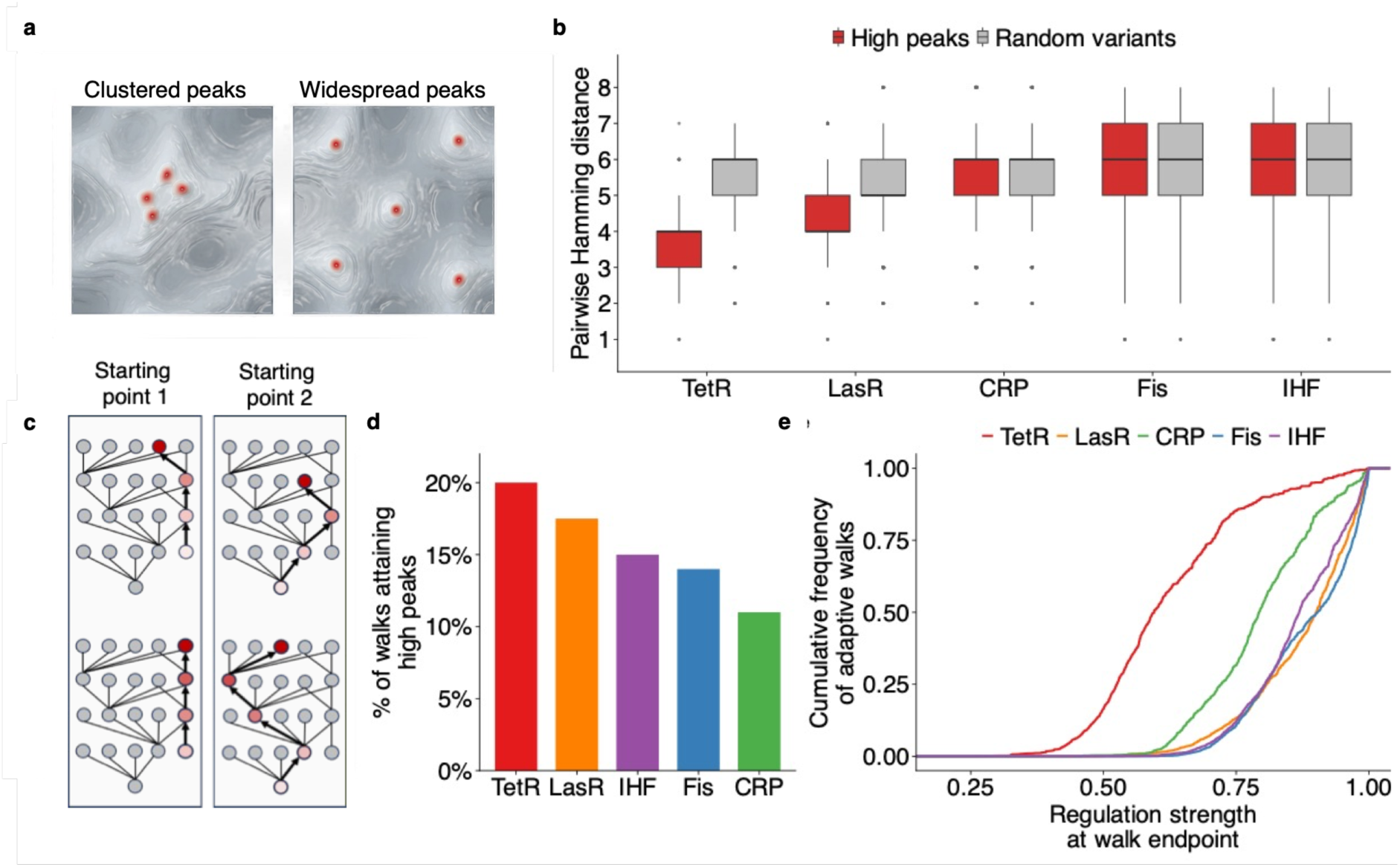
High peaks are non-randomly distributed in genotype space and differentially accessible by adaptive walks. a Schematic of peak classification (Supplementary Methods Section 7). A peak is a genotype whose single-nucleotide neighbours all have lower regulation strength than itself; a high peak is a peak whose regulation strength exceeds that of the wild-type sequence. The five landscapes contain 58 (TetR), 82 (LasR), 61 (CRP), 172 (Fis), and 199 (IHF) high peaks; total numbers of peaks (high + low) are 2,160, 2,581, 2,154, 2,327, and 2,453, respectively. Red points indicate high peaks. **b Pairwise Hamming distances between high peaks (red) and between random non-peak genotypes (grey, sampled from the same landscape and matched in number to the high peaks of the respective TF).** Box plots show the median, interquartile range, and whiskers extending to 1.5× the interquartile range; points indicate outliers. Pairwise distances were calculated among all pairs of high peaks for each landscape, yielding 1,653 pairwise distances for TetR, 3,321 for LasR, 1,830 for CRP, 14,706 for Fis, and 19,701 for IHF. The corresponding numbers of high peaks are 58 (TetR), 82 (LasR), 61 (CRP), 172 (Fis), and 199 (IHF). The number of random genotypes chosen to compute pairwise Hamming distances matches the number of high peaks for each TF. **c. Schematic of adaptive walks from two different starting genotypes.** Each node represents a genotype, and edges connect genotypes that differ by a single mutation. Node colour indicates regulation strength, with darker red indicating higher regulation strength. Arrows indicate specific mutational steps taken during an adaptive walk. Depending on the starting genotype, walks can reach different peaks. **d Percentage of adaptive walks reaching a high peak in each landscape, at N = 10⁸.** Each bar summarises 10^8^ adaptive walks per landscape, corresponding to 10,000 starting genotypes and 10,000 walks per starting genotype. **e. Cumulative distribution function (CDF) of regulation strength at the endpoint of adaptive walks at population size N = 10⁸ (Supplementary Methods Section 9).** Each curve summarises 10⁸ walks per landscape (10,000 starting genotypes × 10,000 walks). Adaptive walks on the TetR landscape (red) reach lower endpoints than walks on global-regulator landscapes (CRP, Fis, IHF), which reach a regulation strength close to the maximum.

**Table 1.**
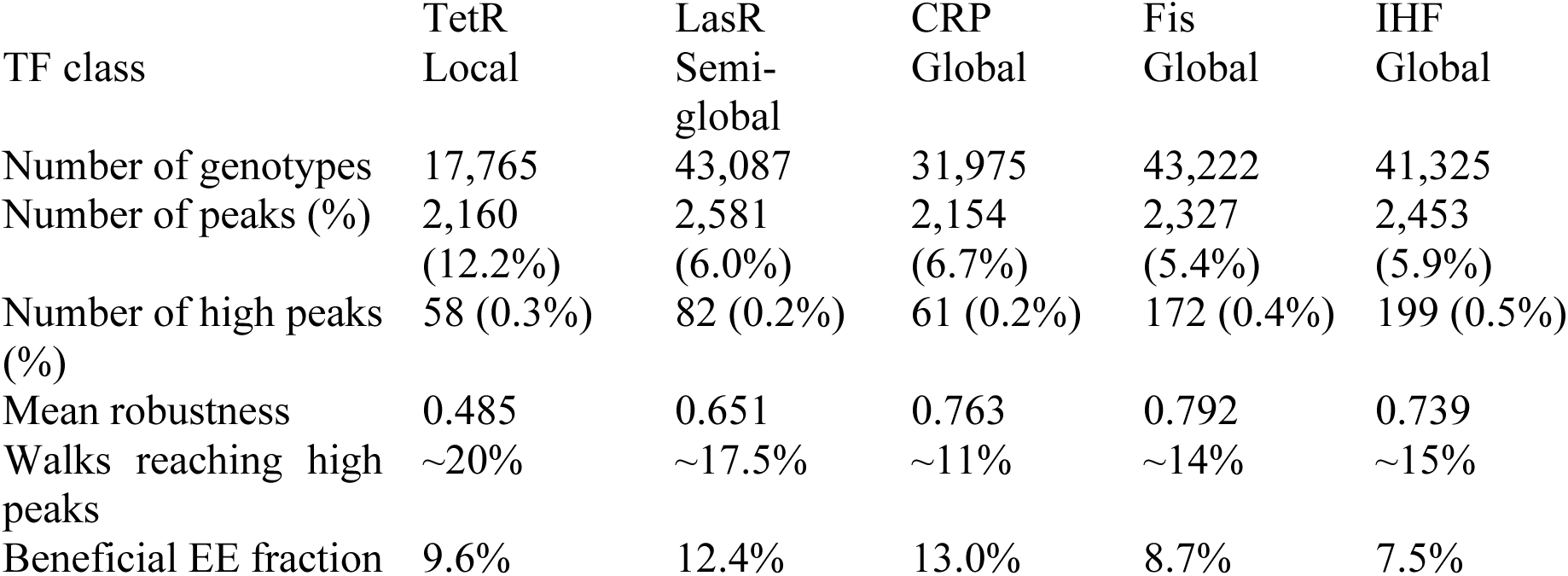
Summary of landscape properties across TFs.

The distribution of high peaks in sequence space depends on the TF class. In the TetR landscape, high peaks are genetically clustered: their median pairwise Hamming distance is substantially smaller than that between randomly sampled genotypes (Wilcoxon rank-sum test, *P* < 2.2 × 10⁻¹⁶; **Fig. 3b; Supplementary Methods Section 8**). In contrast, high peaks in the global landscapes (CRP, Fis, IHF) are more widely dispersed, with distances close to the random expectation. High peaks in the LasR landscape showed an intermediate pattern. In other words alternative strong-binding sites are separated by fewer mutations in the landscape of the local TF than in the landscapes of the global TFs.

### Peak accessibility decreases with regulatory scope

To determine whether these topographic differences affect evolutionary dynamics, we simulated the adaptive evolution of populations, using adaptive walks and the Kimura fixation probability framework ^64–66^ in the strong-selection, weak-mutation regime ^42,43^ (**Fig. 3c; Supplementary Methods Section 9**). We used an effective population size of *N* = 10⁸ individuals, which is representative of natural *E. coli* populations ^67^, and incorporated experimentally determined mutation biases into our walks ^68,69^. For each landscape, we performed 10⁸ adaptive walks (10,000 starting genotypes × 10,000 walks each), each of up to 10 mutational steps. Most walks terminated within 3–5 steps at a local peak (**Supplementary Fig. S6**), indicating that our upper step limit does not bias accessibility estimates.

We define peak accessibility as the fraction of adaptive walks that reach a high peak, that is, a peak whose regulation strength exceeds that of the wild-type TFBS. Observed peak accessibility exceeded that of randomised landscapes for all TFs, where randomisation permuted regulation-strength values across genotypes while preserving network connectivity (all P < 0.001, based on permutation tests with 1,000 randomised landscapes per TF; null hypothesis: observed accessibility is no greater than expected under random assignment of regulation strength to genotypes). Thus, empirical landscape structure facilitates access to high peaks beyond random expectations.

Observed peak accessibility differs systematically among TFs (**Fig. 3d,e**). In the TetR landscape, approximately 20% of walks reach a high peak. Accessibility is slightly lower in the LasR landscape, where approximately 17.5% of walks reach a high peak, and lower still in the global landscapes, i.e., approximately 15% for IHF, 14% for Fis, and 11% for CRP. Because high peaks are defined relative to each TF’s wild-type regulation strength, higher peak accessibility does not necessarily imply higher absolute endpoint regulation strength across landscapes. The distribution of endpoint regulation-strength therefore provides a complementary view of adaptive-walk outcomes (**Fig. 3e**):

### Evolvability-enhancing mutations are abundant and improve adaptive walks

So far, we have shown that mutational robustness, peak distance, and peak accessibility vary with regulatory scope. We next asked whether a fourth aspect of landscape organization and evolutionary potential also varies with scope: the prevalence and evolutionary impact of evolvability-enhancing (EE) mutations. In fitness landscapes, EE mutations are mutations that create genetic backgrounds in which subsequent mutations are more likely to be beneficial than expected under additivity — not simply because the EE mutation itself is beneficial, but because it increases the mean fitness of neighbouring genotypes relative to the additive expectation ^49^. We apply this framework to regulation-strength landscapes under the assumption that regulation strength is a fitness proxy. Thus, in our analysis, an EE mutation is one that increases the mean regulation strength of neighbouring genotypes beyond what is expected from its own direct effect on regulation strength (**Fig. 4a**; **Supplementary Methods Section 14.1**). Such mutations can contribute to adaptive evolution through their own direct effects while also altering the set of beneficial mutations available subsequently, without requiring second-order selection on evolvability ^49^.

**Figure 4.**
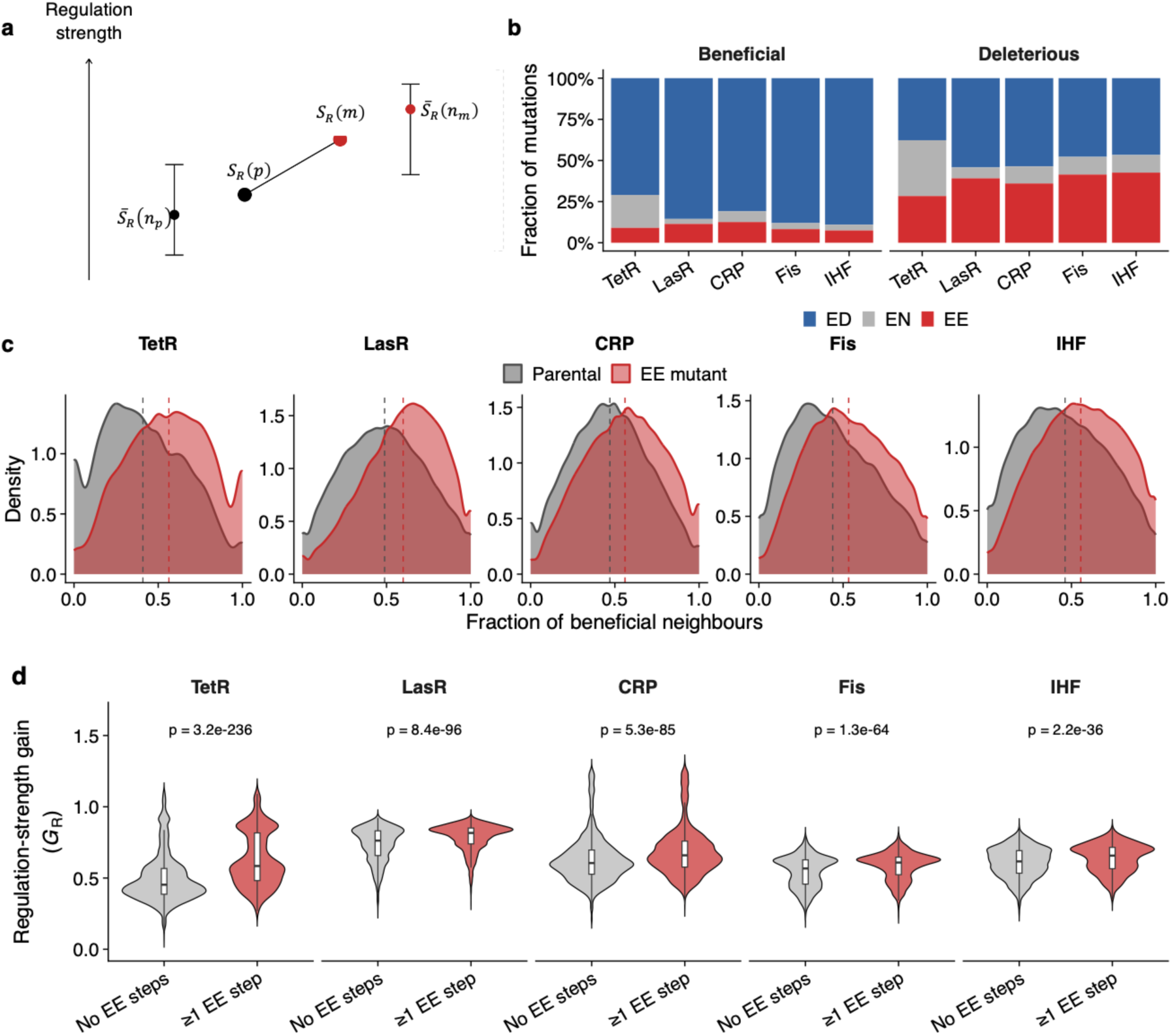
**Evolvability-enhancing mutations increase regulation-strength gains during adaptive walks. a. Schematic of the EE mutation framework, adapted from ref.**^49^. For a mutation from a parental genotype *p*to a single-mutant genotype *m*, the direct effect on regulation strength is Δ*S_R_* = *S_R_*(*m*) − *S_R_*(*p*). A mutation is classified as evolvability-enhancing (EE) if the mean regulation strength of all genotypes in the mutant neighbourhood, *S̄_R_* (*n_m_*), exceeds the additive expectation based on the parental neighbourhood, *S̄_R_* (*n_p_*), and the mutation’s direct effect. Mutations not satisfying this criterion are classified as evolvability-neutral (EN) or evolvability-reducing (ED) according to the sign of the neighbourhood effect. **b**. **Fraction of mutations classified as EE (red), EN (grey), or ED (blue) among beneficial (left) and deleterious (right) mutations for each landscape.** The percentages of EE mutations among beneficial mutations are 9.6% (TetR), 12.4% (LasR), 13.0% (CRP), 8.7% (Fis), and 7.5% (IHF); among deleterious mutations they are 27.4%, 39.8%, 36.5%, 41.6%, and 42.0%, respectively. **c**. **Distribution of the fraction of beneficial neighbours for each EE mutant genotype (red) and its parental genotype, i.e., before the EE mutation (grey).** The vertical axis shows probability density, and vertical dashed lines indicate medians. Median fractions of beneficial neighbours for parental genotypes versus EE mutants are 0.40 vs. 0.57 (TetR), 0.50 vs. 0.62 (LasR), 0.47 vs. 0.57 (CRP), 0.41 vs. 0.53 (Fis), and 0.44 vs. 0.56 (IHF). Paired Wilcoxon signed-rank tests reject the null hypothesis that parental genotypes and EE mutants have the same median fraction of beneficial neighbours (all P < 2.2 × 10⁻¹⁶; number of genotype pairs = 5,441 for TetR, 42,206 for LasR, 28,713 for CRP, 30,986 for Fis, and 25,715 for IHF). **d**. **Final regulation-strength gain of adaptive walks containing no EE steps (grey) versus walks containing at least one EE step (red), for each landscape at N = 10^8^.** Regulation-strength gain (*G_R_*) is defined as 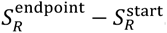 Box plots inside the violins show the interquartile range, with the median marked by a horizontal line and whiskers extending to 1.5× the interquartile range. P-values are derived from two-sided Mann–Whitney U tests rejecting the null hypothesis that final regulation-strength gain is drawn from the same distribution for walks with and without EE mutstions. Cliff’s δ effect sizes are +0.43 (TetR), +0.25 (LasR), +0.23 (CRP), +0.22 (Fis), and +0.17 (IHF), where positive values indicate larger regulation-strength gains for EE-containing walks. To reduce non-independence among walks sharing the same starting genotype, this comparison is based on one randomly selected walk per starting genotype, yielding 10,000 walks per landscape partitioned by EE-step inclusion: TetR (n_no-EE = 7,355, n_EE = 2,645), LasR (6,656; 3,344), CRP (5,468; 4,532), Fis (7,182; 2,818), and IHF (7,558; 2,442). The largest EE-associated benefit, an approximately 27% increase in mean regulation-strength gain, occurs in the TetR landscape.

We classified all single-nucleotide mutations as evolvability-enhancing (EE), evolvability-neutral (EN), or evolvability-reducing (ED), separately for beneficial and deleterious mutations. All five landscapes harbour EE mutations (**Fig. 4b**; **Supplementary Figs. S7, S8**). Among beneficial mutations, the fraction classified as EE ranges from 7.5% in IHF to 13.0% in CRP; among deleterious mutations, it ranges from 27.4% in TetR to 42.0% in IHF. These fractions exceed those observed in randomised landscapes, in which regulation-strength values were permuted across genotypes while preserving genotype-network structure (P< 0.05; permutation tests for 1,000 randomised landscapes per TF; null hypothesis: observed EE prevalence is no greater than expected under random assignment of regulation strength to genotypes; **Supplementary Figs. S7, S8**).

EE mutations also clearly affect the incidence of beneficial mutations. That is, in all landscapes, the genotypes they produce have a higher fraction of beneficial neighbours than pre-mutation genotypes (median fraction of beneficial neighbours, pre-mutation vs. EE mutant: TetR 0.40 vs. 0.57; LasR 0.50 vs. 0.62; CRP 0.47 vs. 0.57; Fis 0.41 vs. 0.53; IHF 0.44 vs. 0.56; paired Wilcoxon signed-rank tests rejecting the null hypothesis that EE mutants and their parental genotypes have the same median fraction of beneficial neighbours, all P < 2.2 × 10⁻¹⁶; number of genotype pairs: 5,441 for TetR, 42,206 for LasR, 28,713 for CRP, 30,986 for Fis, and 25,715 for IHF; **Fig. 4c**). Thus, EE mutations expand the set of beneficial mutational steps available to adaptive evolution. Because this effect may propagate to subsequent mutations, even rare EE mutations may substantially shift the trajectory of an adaptive walk, a prediction we tested next.

Adaptive walks that include at least one EE mutation entail greater gains in regulation-strength than walks composed entirely of non-EE mutations (two-sided Mann–Whitney U tests rejecting the null hypothesis that regulation-strength gains are drawn from the same distribution for walks with and without EE mutations: P = 3.2 × 10⁻²³⁶ for TetR, 8.4 × 10⁻⁹⁶ for LasR, 5.3 × 10⁻⁸⁵ for CRP, 1.3 × 10⁻⁶⁴ for Fis, and 2.2 × 10⁻³⁶ for IHF; Cliff’s δ = +0.43, +0.25, +0.23, +0.22, and +0.17, respectively; **Fig. 4d**; per-walk trajectories in **Supplementary Fig. S10**). This influence of EE mutations is largest in the TetR landscape, where EE-containing walks show an approximately 27% larger mean gain in regulation strength than walks without EE mutations. It persists across three population sizes spanning six orders of magnitude (N = 10²– 10⁸; **Supplementary Fig. S12**), indicating that it is not restricted to populations in which genetic drift is very weak.

The relationship between robustness and EE mutations is less consistent among TFs. Within landscapes, more robust genotypes have fewer EE mutants in TetR, LasR, and CRP (Spearman ρ = −0.066 to −0.107; all *P* < 10⁻¹⁴), but the opposite holds for Fis (ρ = +0.046, P = 4.9 × 10⁻²², n = 43,210) and IHF (ρ = +0.013, P = 0.01, n = 41,296) (**Supplementary Fig. S11**). Across landscapes, EE prevalence and mean robustness are uncorrelated (Spearman ρ = −0.10, *P* = 0.95; *n* = 5; **Supplementary Fig. S9**).

## Discussion

Our results show that TFBS adaptive landscapes properties vary systematically with the regulatory scope of their cognate TFs. Specifically, four properties — mutational robustness, high-peak distribution, peak accessibility, and the within-adaptive-walk benefit of evolvability-enhancing (EE) mutations — vary along a local-to-global axis (Fig. 5). The local TetR landscape sits at one extreme: its genotypes have the lowest mean mutational robustness, its high peaks are the most genetically clustered, its high peaks are evolutionarily most accessible during adaptive walks, and EE-containing adaptive walks cause the largest gain in regulation strength. The global landscapes of CRP, Fis, and IHF show the opposite pattern: their genotypes are more robust to mutation, their high peaks are more dispersed, high peaks are less accessible, and the benefit of EE mutations during adaptive walks is smaller. The semi-global LasR landscape has intermediate features. This pattern indicates that regulatory scope is associated with the topography of TFBS adaptive landscapes, although the global landscapes are not strictly ordered across all metrics we considered.

**Figure 5.**
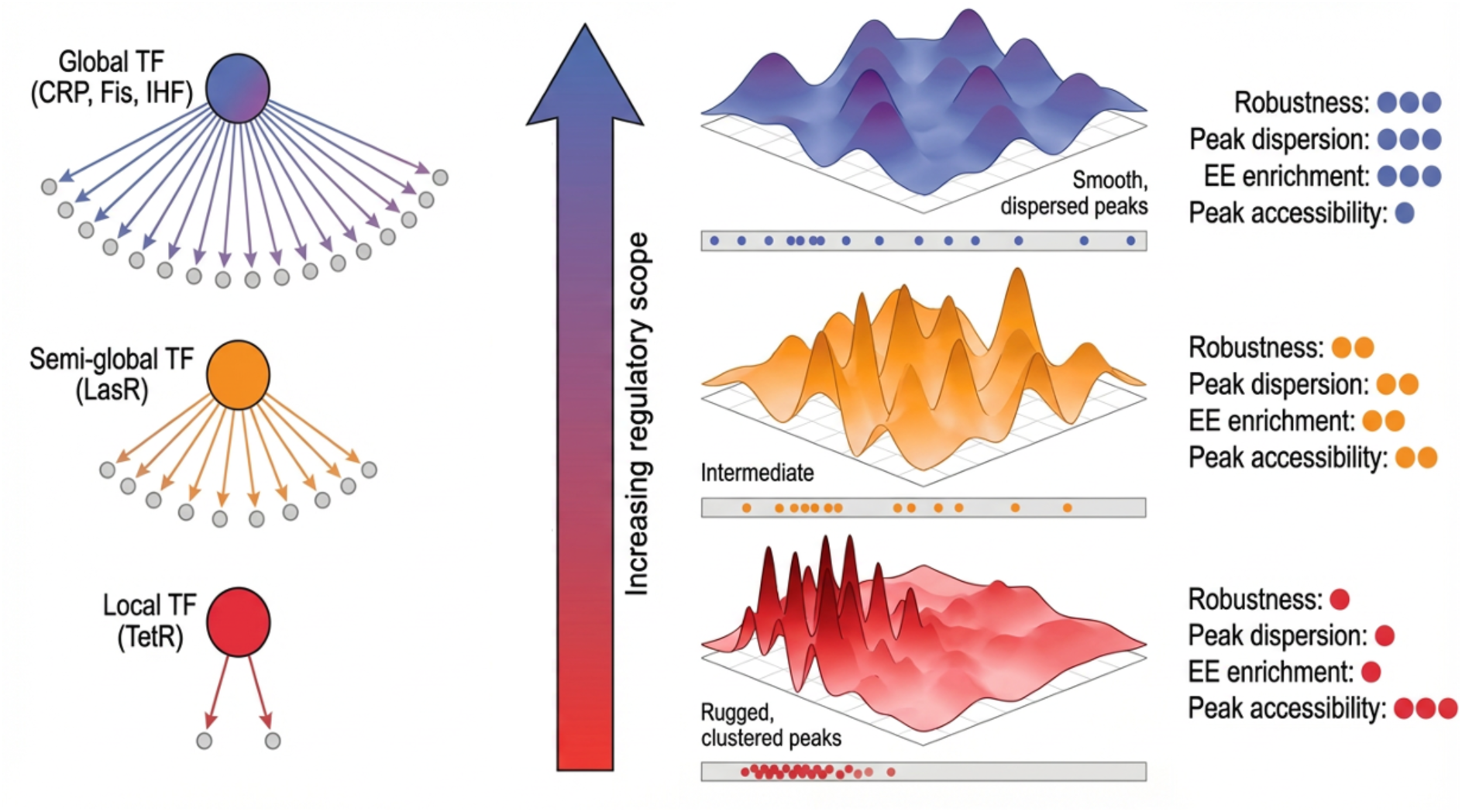
Summary of landscape properties across regulatory scope. Schematic showing transcription factors within a gene regulatory network hierarchy (see also Fig. 1a). The local regulator (bottom) controls few genes. Its TFBS landscape has low mean mutational robustness, genetically clustered high peaks, high peak accessibility, and the largest within-walk benefit of evolvability-enhancing (EE) mutations. Global regulators (top) control many genes. Their TFBS landscapes have higher mean mutational robustness, more dispersed high peaks, lower peak accessibility, and smaller within-walk EE benefits. The semi-global regulator (middle) has intermediate features. The gradient (thick vertical arrow) reflects the systematic relationship between regulatory scope and TFBS landscape topography.

The landscape properties of the local TetR regulator are consistent with the demands of tight and specific regulation. TetR represses the *tetA* efflux pump. Its leaky expression imposes fitness costs by draining resources and perturbing membrane potential ^70–72^. Similar constraints apply to other local repressors such as LacI ^31,73,74^, where repression strength must be tightly maintained ^8,9,75^. In this context, the clustering and high accessibility of high-regulation-strength peaks may facilitate recovery of strong repression after mutations that weaken regulation. However, the same landscape also has low mean mutational robustness: single mutations tend to have large effects on regulation strength. Rather than a trade-off, we interpret low robustness and high peak accessibility as joint consequences of the high binding specificity of a local repressor. A sharp, information-rich motif makes regulation strength fall steeply as sequences depart from the consensus, so mutations have large effects (low robustness)^57,76^; the same specificity concentrates strong-binding sequences into a dense cluster in sequence space, so that high-regulation-strength peaks lie close together and are readily reached by adaptive walks (high accessibility)^77,78^. Sensitivity to mutation and ready access to strong-regulation genotypes are thus two facets of the same specific, steep regulatory landscape, rather than properties in tension.

The global landscapes show a different organisation. CRP, Fis, and IHF regulate hundreds of genes and contribute to chromosome organisation^8,9,79^. Their biological roles rely on distributed binding across many genomic sites rather than regulation through one or a few highly specific sites. The higher mean robustness of genotypes in these landscapes indicates that single mutations usually produce smaller changes in regulation strength. This may be compatible with global regulation, where many genomic binding sites must remain functional despite sequence variation. The wider dispersion of high peaks indicates that alternative strong-regulation genotypes are separated by more mutations, and their lower accessibility suggests that adaptive walks under directional selection for increased regulation strength reach these high peaks less often. Together with the broad distribution of predicted regulation strengths across genomic CRP, Fis, and IHF binding sites (**Supplementary Fig. S2**), these observations suggest that maximal regulation strength may not be the typical evolutionary endpoint for global TFBSs. Instead, adaptive evolution of such TFBSs may often involve tuning sites within a broader range of moderate-to-high regulation strengths ^80–82^. However, the position of IHF on the robustness axis also shows that regulatory scope is not the only determinant of landscape topography. Differences in TF biology, such as ligand dependence, DNA-bending activity, dimerisation mode, and the balance between direct transcriptional control and chromosome organisation, may further shape TFBS landscape structure ^57,83^.

The LasR landscape occupies an intermediate position between the local TetR landscape and the global landscapes. In *P. aeruginosa*, LasR regulates quorum-sensing genes in response to the autoinducer 3-oxo-C12-homoserine lactone, coupling gene expression to cell-density-dependent signal concentration^18,19,84^. Its TFBS landscape has intermediate mutational robustness, high-peak clustering, and high-peak accessibility. This position is consistent with the biology of LasR as a regulator that directly binds fewer genomic regions than the global regulators analysed here, but controls a broader regulon than TetR through the quorum-sensing hierarchy. In evolutionary terms, this intermediate landscape topography allows LasR binding sites to tolerate more mutations than TetR binding sites without abolishing regulation, while still retaining access to high-regulation-strength genotypes.

Evolvability-enhancing mutations, previously described in RNA and protein fitness landscapes ^49^, are also present in all TFBS landscapes analysed here. In these landscapes, EE mutations increase the fraction of beneficial neighbouring genotypes and thereby expand the set of adaptive mutations available after an EE mutation. Adaptive walks containing at least one EE mutation reach higher regulation-strength gains than walks without EE mutations in all five landscapes, an effect that is largest in the TetR landscape. This advantage persists for effective population ranging from N = 10^2^ to N = 10^8^, indicating that the effect is not restricted to large populations in which genetic drift is negligible. Thus, EE mutations appear to be a general feature of TFBS adaptive landscapes and can alter adaptive outcomes under selection for increased regulation strength.

Several caveats qualify these conclusions. First, we analyse TFBS landscapes, not TF evolution or organismal fitness landscapes. Our adaptive walks use regulation strength as a proxy for fitness under the assumption that stronger regulation by the cognate TF is favoured. This assumption allows us to compare landscape topography across TFs, but it does not imply that stronger regulation is always beneficial in nature. Natural selection may favour intermediate regulation strengths, context-dependent expression, or reduced non-cognate binding. Second, the landscape differences we observe may result from TF–DNA biophysics, from the evolutionary history of each regulator, or from both. For example, highly specific TF–DNA recognition may naturally produce clustered sets of strong-binding sequences, whereas more promiscuous recognition may distribute strong-regulation genotypes more broadly across sequence space. Distinguishing intrinsic biophysical effects from historical or selective effects will require larger comparative datasets across TF families and regulatory scopes.

A third caveat is that our observations derive from bacterial TFBS landscapes and may not transfer directly to eukaryotic regulatory systems. Bacterial regulatory regions often contain few TFBSs, so changes in individual binding sites can have large effects on gene expression^44,85–87^. In eukaryotes, by contrast, regulatory specificity often emerges from combinations of multiple low-information binding sites within enhancers ^27,78^ ^88^ ^44,76,85^, and mutational robustness may arise at the level of enhancer architecture rather than individual TFBSs ^45,47,89,90^. Extending similar landscape measurements to multi-site regulatory regions will therefore be important for testing whether the relationship between regulatory scope and TFBS landscape structure generalises beyond bacteria.

Future work should extend these analyses to more TFs, multiple binding sites within the same regulatory region, and landscapes measured under different environmental conditions. This broader sampling will be essential to test whether the local-to-global gradient observed here holds across TF families and to determine whether relationships among robustness, peak accessibility, and EE mutations are general features of TFBS evolution. Experimental evolution systems in which TFs are recruited to new regulatory targets ^12^ could provide a direct test of whether measured TFBS landscape properties predict regulatory rewiring. More broadly, our results suggest that the evolutionary potential of cis-regulatory sequences depends not only on their local sequence properties, but also on the regulatory scope of the TFs that bind them.

## Methods

### Overview

This study analyses Sort-Seq datasets for the five bacterial transcription factors TetR, LasR, CRP, Fis, and IHF. We generated the TetR, CRP, Fis, and IHF datasets in previous studies using the same experimental and computational framework^29,30^. We generated the LasR dataset specifically for this study, following the same standardised Sort-Seq protocol and analysis pipeline (**Supplementary Methods Section 2**). Briefly, each library consisted of TFBS variants cloned upstream of a GFP reporter gene and regulated by the cognate TF. Cells carrying these libraries were sorted into 13 fluorescence bins by fluorescence-activated cell sorting, and variants in each bin were quantified by deep sequencing. The distribution of each variant across bins was then used to infer its expression level and regulation strength.

### Calculating regulation strength

For each TFBS variant, we estimated GFP expression *e* as the weighted average of the fluorescence bins in which the variant was observed, i.e.,

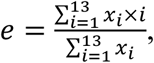

where *x_i_* is the number of sequencing reads observed for that variant in fluorescence bin *i*, and (*i* = 1, … ,13) indexes bins from lowest to highest fluorescence. Because higher GFP expression corresponds to weaker TF-mediated regulation in our assay, we inverted this expression scale to obtain raw regulation strength:

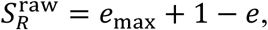

where *e*_max_ = 13. We then normalised raw regulation strength within each landscape as

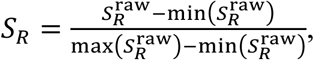

yielding a normalised regulation-strength score *S_R_* that ranges from zero for the weakest observed regulation to one for the strongest observed regulation.

### Combining replicates

Each dataset consists of data from three biological replicate Sort-Seq experiments for TetR, CRP, Fis, and IHF, and from four biological replicate experiments for LasR. We retained a TFBS variant for downstream analysis only if it was detected in all biological replicates, had at least 30 total sequencing reads across all fluorescence bins in each replicate, and had a coefficient of variation in regulation strength below 0.5 across replicates (**Supplementary Methods Section 4**). For each retained variant, we used the mean normalised regulation strength across replicates for all downstream analyses.

### Genotype networks

For each TF, we represented the TFBS landscape as a genotype network using the Python package igraph^91^. Nodes correspond to TFBS variants, and edges connect pairs of variants that differ by a single nucleotide. For analyses of adaptive walks, we represented each edge as directed from the lower to the higher regulation strength variant. Because incomplete sampling can fragment genotype networks, we restricted all analyses to the largest weakly connected component of each landscape^54^, which comprises approximately 97% of all retained genotypes for each TF (**Supplementary Methods Section 5**).

### Distribution of mutational effects

For each single-nucleotide mutation connecting two neighbouring genotypes, we computed its effect on regulation strength as *ΔS_R_* = *S_R_*(*j*) − *S_R_*(*i*), where *i* is the pre-mutation genotype and *j* is the mutant genotype. We analysed the distribution of mutational effects across each landscape and within three regulation-strength classes (low, medium, and high), defined by tertiles of *S_R_* within each landscape (**Supplementary Methods Section 7.3**). We compared distributions of mutational effects between pairs of landscapes using two-sample Kolmogorov– Smirnov tests. The null hypothesis was that the mutational effects in each pair of landscapes were drawn from the same distribution (**Supplementary Methods Section 11**).

### Mutational robustness

We quantified the sensitivity of each genotype *g* to mutation as

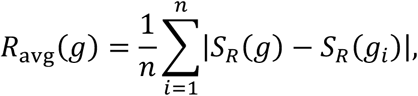

where *g_i_* denotes one of the *n* single-mutant neighbours of genotype *g*, and *S_R_*(g) denotes the normalised regulation strength of genotype *g*. We report mutational robustness as

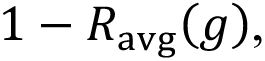

so that higher values indicate greater robustness (**Supplementary Methods Section 6.1**). To compare the robustness of peak and non-peak genotypes within each landscape, we used two-sided Wilcoxon rank-sum tests. Because our large sample sizes yield vanishingly small *P* values even for negligible differences, we additionally report Cliff’s delta (δ) as a sample-size-independent effect size. For two groups with values *xᵢ* and *yⱼ* of sizes n_x_ and n_y_, *δ* = [(*x*; > *yⱼ*) − (*x*; < *yⱼ*)]/(*n_x_n_y_*). It ranges from −1 to +1, equals 0 when the two distributions overlap completely, and is positive (negative) when values in the first group tend to exceed (fall below) those in the second; its magnitude measures the degree of stochastic dominance. δ is a linear rescaling of the Mann–Whitney *U* statistic *δ* = 2*U*/(*n_x_n_y_*) − 1 and thus reflects the same pairwise comparisons as the rank-sum test. We interpret |δ| as negligible (<0.147), small (0.147–0.33), medium (0.33–0.474), or large (≥0.474)^59^.

To test whether robustness differed among landscapes, we used a Kruskal–Wallis test^60^, with the null hypothesis that robustness values are drawn from the same distribution across landscapes. We then performed pairwise Dunn’s post-hoc tests^61^ with Bonferroni correction to identify which pairs of landscapes differed in mutational robustness.

### Peaks and pairwise distances

A peak is a genotype whose single-nucleotide neighbours all have lower regulation strength than the genotype itself (**Supplementary Methods Section 7.1**). We defined a high peak in each landscape as a peak whose regulation strength exceeds that of the wild-type TFBS for the corresponding TF (**Supplementary Methods Section 7.2**). To quantify the spatial distribution of high peaks, we calculated pairwise Hamming distances among all high peaks within each landscape. We compared these distances with pairwise Hamming distances among an equal number of randomly sampled non-peak genotypes from the same landscape (**Supplementary Methods Section 8**). We used Wilcoxon rank-sum tests to test the null hypothesis that pairwise distances among high peaks and pairwise distances among randomly sampled non-peak genotypes are drawn from the same distribution.

### Simulated adaptive walks

We simulated adaptive walks under the strong-selection, weak-mutation regime (**Supplementary Methods Section 9.1**). In this regime, populations are assumed to be monomorphic most of the time, and adaptation proceeds through the fixation of beneficial single mutations. For a mutation from genotype *i* to neighbouring genotype *j*, we calculated the selection coefficient as

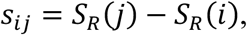

where *S_R_* is normalised regulation strength. We considered mutations with *s_i_*_j_ > 0 as beneficial, and calculated their fixation probability using Kimura’s formula^65^,

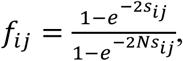

where *N* is the effective population size. For most simulations, we used N = 10^8^, and assessed sensitivity of our results to population size using N = 10^5^ and N = 10^2^. We incorporated experimentally determined mutation biases for *E. coli* into the transition probabilities of adaptive walks ^68,69^. For each landscape, we randomly selected 10,000 non-peak starting genotypes and initiated 10,000 walks from each starting genotype, yielding 10^8^ walks per landscape. Walks were allowed to proceed for up to 10 mutation-fixation steps.

### Randomised landscapes

We generated 1,000 randomised landscapes per TF by permuting regulation-strength values across genotypes while preserving genotype-network connectivity (**Supplementary Methods Section 10**). This procedure preserves the structure of sequence space and the empirical distribution of regulation strengths, but randomises the association between genotype and regulation strength. We used these randomised landscapes to test whether observed landscape properties exceeded expectations under random assignment of regulation strength to genotypes. For each landscape statistic we studied, the null hypothesis was that the observed value was no greater than expected from the corresponding randomised landscapes. We calculated P-values and bootstrap 95% confidence intervals based on the 1,000 randomized landscapes.

### EE mutation classification

We classified single-nucleotide mutations as evolvability-enhancing (EE), evolvability-neutral (EN), or evolvability-reducing (ED), following ref. ^49^ (**Supplementary Methods Section 14**). For each mutation from a parental genotype *p* to a mutant genotype *m*, we first calculated the mutation’s direct effect on regulation strength as

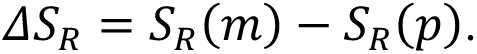

We then compared the mean regulation strength of the single-mutant neighbourhood of *m*, *S̅_R_*(*n_m_*), with that of the single-mutant neighbourhood of *p*, *S̅_R_*(*n_p_*). For beneficial mutations, additivity predicts that the mutant neighbourhood should differ from the parental neighbourhood by the mutation’s direct effect, *ΔS_R_*. For deleterious mutations, we used the stricter criterion from ref. ^49^, requiring the mutant neighbourhood to have higher mean regulation strength than the parental neighbourhood to be classified as EE. We therefore evaluated the neighbourhood effect as

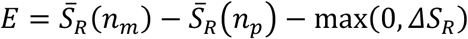

We classified a mutation as EE if E > 0, as ED if E < 0, and as EN if E did not differ significantly from zero. For each mutation, we tested the null hypothesis that E = 0 with a two-sided one-sample *t*-test, comparing the neighbourhood-mean difference S̄*_R_*(*n_m_*) − *S_R_*(*n_p_*)against its null expectation *max*(0, *ΔS_R_*). We obtained the standard error of this difference by propagating the per-genotype standard errors of regulation strength, estimated across replicates: for a neighbourhood of *n* genotypes with individual standard errors *SE_i_*, the standard error of its mean is (1/*n*) · √(Z*_i_SE_i_*²)(**Supplementary Methods Section 14**). We corrected the resulting *P*-values across all mutations using the Benjamini–Hochberg procedure at a false discovery rate of 1%. We assessed whether EE mutations were enriched relative to random expectation using the randomised landscapes described in the previous section. We also assessed the consequences of EE mutations for adaptive walks by comparing regulation-strength gain, walk length, and the fraction of walks terminating at low peaks between walks containing at least one EE mutation and walks containing no EE mutations (**Supplementary Methods Section 14.8**).

## Data Availability

All code, processed data, plasmid sequences, and primer sequences are available in our Zenodo repository (10.5281/zenodo.21531555), which will be made publicly accessible upon acceptance of the manuscript. Sequencing data are deposited in the Sequence Read Archive under BioProject PRJNA1499622 (LasR, generated for this study), PRJNA1162449 (CRP, Fis, and IHF) and PRJNA1019339 (TetR). The latter two datasets were generated in our previous studies.

## Acknowledgements

We would like to acknowledge financial support by Swiss National Science Foundation grant 310030_208174. We would also like to thank the UZH Cytometry Facility for technical support.

## Author Contributions

C.A.W. and A.W. conceived the study and designed the experiments. C.A.W. executed the LasR Sort-Seq experiments, performed all computational analyses, and generated all figures. C.A.W. and A.W. wrote the manuscript.

## Competing Interests

The authors declare no competing interests.

See **Supplementary Methods Section 4** for full data filtering summary

## Notes

### Competing Interest Statement

The authors have declared no competing interest.

