## Supporting Information for "Regulatory scope shapes the adaptive landscapes of bacterial transcription factor binding sites"

**Supplementary Methods**

**1. Sort-Seq experimental overview**

**2. LasR Sort-Seq experiment**

- **2.1. Strain and plasmid construction**
- **2.2. Library design and cloning**
- **2.3. Sort-Seq and FACS**

**3. Calculating regulation strengths**

**4. Combining biological replicates**

**5. Genotype network construction**

**6. Mutational robustness**

- **6.1. Mean absolute difference ($R_{avg}$)**
- **6.2. Choice of neighbourhood radius**

**7. Classification of genotypes**

- **7.1. Peaks and plateaus**
- **7.2. High peaks**
- **7.3. Regulation strength classes**

**8. Pairwise genetic distances between high peaks**

**9. Simulated adaptive walks**

- **9.1. Model**
- **9.2. Mutation bias**
- **9.3. Simulation procedure**
- **9.4. Population sizes**
- **9.5. Accessibility metrics**

**10. Randomised null model for accessibility**

**11. Distribution of mutational effects (DFE) analysis**

**12. Statistical comparisons across TF classes**

- **12.1. Robustness across TF classes**

12.2. Peaks versus non-peaks

12.3. High-peak distance comparisons

13. Position weight matrix (PWM) analysis

14. Evolvability-enhancing (EE) mutation classification

14.1. Framework

14.2. Statistical test

14.3. Measurement variance

14.4. Multiple testing correction

14.5. Randomised null model for EE enrichment

14.6. Coverage

14.7. Validation

14.8. Consequences for adaptive walks

14.9. Fraction of beneficial neighbours and adjusted neighbourhood mean

14.10. Within-landscape coupling between robustness and EE prevalence

14.11. Cross-landscape relationship between robustness and EE prevalence

15. Software and computational environment

**Supplementary Figures**

S1. Distribution of regulation strengths across landscapes

S2. Distribution of predicted regulation strengths for genomic TFBSs

S3. Relationship between the number of neighbours and regulation strength across TFBS landscapes

S4. Distribution of mutational effects (DFE) by regulation-strength class

S5. Peaks are less mutationally robust than non-peak genotypes in every landscape

S6. Adaptive walks settle quickly: most reach a local peak within 3–5 mutational steps

S7. EE mutants have higher mean neighbour regulation strength than their parental genotypes

S8. EE mutants increase adjusted mean neighbour regulation strength.

**S9. Cross-landscape relationship between mean robustness and EE prevalence**

**S10. Adaptive walks containing EE steps reach higher endpoints across all five landscapes**

**S11. Within-landscape coupling between robustness and EE prevalence is weak and inconsistent in sign**

**S12. Walks with EE mutations show larger regulation-strength gains at different population sizes**

### Supplementary Methods

#### 1. Sort-Seq experimental overview

This study analyses Sort-Seq datasets for five bacterial transcription factors: TetR (local regulator), LasR (semi-global regulator), as well as CRP, Fis, and IHF (global regulators). We generated the datasets for TetR, CRP, Fis, and IHF and described them in detail in previous publications[1,2] that include TF-specific considerations such as strain backgrounds, induction conditions, and gating strategies. We generated the LasR dataset for this study (**Section 2**). We processed all five datasets with the same computational pipeline described below.

#### 2. LasR Sort-Seq experiment

##### 2.1 Strain and plasmid construction

We performed all cloning and Sort-Seq experiments for LasR in *Escherichia coli* strain DH5α (SIG10-MAX, Sigma Aldrich, CMC0004), which we chose for its high transformation efficiency and stable plasmid propagation. We derived the LasR expression plasmid from the modular backbone pCAW-Sort-Seq-V2[1], which contains a pBBR1 origin of replication, a chloramphenicol resistance cassette, and dual expression modules for a transcription factor and a fluorescent reporter.

We cloned the *lasR* coding sequence upstream of the mScarlet-I fluorescent protein in a bicistronic construct under the control of the aTc-inducible pLtetO-1 promoter. We cloned the corresponding TFBS library upstream of a superfolder GFP (sfGFP) reporter gene, which allows a quantitative measurement of LasR-mediated transcriptional regulation. We performed plasmid assemblies via

Gibson Assembly and sequence-verified them by Sanger sequencing. We transformed final plasmids into SIG10-MAX cells.

#### 2.2 Library design and cloning

We designed the LasR TFBS library based on the LasR consensus binding motif from RegulonDB[3]. We randomized the eight positions with the highest information content, which yields a theoretical diversity of $4^8$ = 65,536 unique sequence variants. The library was synthesised as a single-stranded DNA oligo pool (IDT Ultramers) and converted to double-stranded DNA via PCR with Q5 High-Fidelity DNA Polymerase (NEB), gel-purified (Monarch DNA Gel Extraction Kit, NEB), and ligated upstream of the sfGFP reporter in the LasR plasmid backbone. We electroporated ligation products into SIG10-MAX cells and plated the transformed cells on chloramphenicol LB agar to generate a pooled library with >100-fold coverage.

#### 2.3 Sort-Seq and FACS

We grew transformed cells in LB medium to mid-log phase and induced TF expression with 100 ng/mL anhydrotetracycline (aTc) for 4 hours at 37 °C with shaking. Following induction, we sorted cells using a BD FACSAria III into 13 logarithmically spaced bins based on GFP fluorescence intensity. We restricted sorting to the mScarlet-I-positive population to ensure analysis of cells expressing LasR. We collected sorted cells from each bin into SOC medium, regrew them overnight, and extracted plasmid DNA using the QIAprep Spin Miniprep Kit (Qiagen). We PCR-amplified the TFBS region and sequenced it using Illumina HiSeq (150 bp paired-end mode). We performed four independent biological replicates of these sorting experiments.

**3. Calculating regulation strengths**

In Sort-Seq experiments, sequences may appear in more than one fluorescence bin due to random mis-sorting and stochastic gene expression noise[4–6]. Following previous studies[1,2], we estimated the GFP expression level $e$ associated with each TFBS sequence as the weighted average of the fluorescence bins in which the sequence was observed. For each sequence, we calculated expression as

$$e = \frac{\sum_{i=1}^{13} x_i \times i}{\sum_{i=1}^{13} x_i}$$

where $x_i$ is the number of sequencing reads observed for that sequence in fluorescence bin $i$, and $(i = 1, \ldots, 13)$ indexes bins from lowest to highest GFP fluorescence. This yielded a continuous expression value $e$ between 1 and 13. Because high GFP expression corresponds to weak TF-mediated regulation in our assay, we inverted the expression scale to obtain raw regulation strength,

$$S_R^{\text{raw}} = e_{\text{max}} + 1 - e,$$

where $e_{\text{max}} = 13$. We then normalised raw regulation strength within each landscape as

$$S_R = \frac{S_R^{\text{raw}} - \min(S_R^{\text{raw}})}{\max(S_R^{\text{raw}}) - \min(S_R^{\text{raw}})},$$

which yields a normalised regulation-strength score ($S_R$) ranging from zero for the weakest observed regulation to one for the strongest observed regulation.

### 4. Combining biological replicates

We combined data from independent biological replicates (three replicates for TetR, CRP, Fis, and IHF; four for LasR) as follows:

- **Presence filter**: we retained only sequence variants present in all replicates.
- **Read depth filter**: we removed variants that had fewer than 30 total reads across all 13 bins in at least one replicate.
- **Reproducibility filter**: we computed the coefficient of variation (CV) in regulation strength across replicates for each variant. We excluded variants with CV > 0.5, following standard practice in transcriptional regulation studies using fluorescent reporters, which accept CVs up to this magnitude as admissible due to acceptable transcriptional noise and technical variability[7,8].
- **Averaging**: we computed final regulation strengths as the arithmetic mean across replicates.

As a result of these filtering steps, we retained the following numbers of TFBS variants: TetR: 17,765; LasR: 43,087; CRP: 31,975; Fis: 43,222; IHF: 41,325.

### 5. Genotype network construction

We constructed genotype networks using in-house Python scripts and the python *igraph* package[9] (also used in R for downstream analyses). In these networks, each node represents a TFBS variant, and an edge connects two variants that differ by exactly one nucleotide (Hamming distance = 1). Each edge is directed from the variant with lower regulation strength to the variant with higher regulation strength. We extracted the largest weakly connected component ("giant component"[10]) of each network and used it for all subsequent analyses. In all five landscapes, the giant component comprised ~97% of sequenced genotypes.

### 6. Mutational robustness

#### 6.1 Mean absolute difference in regulation strength ($\boldsymbol{R}_{\mathbf{av}}$)

We quantified the mutational sensitivity of each genotype *g* based on its local mutational neighbourhood, defined as the set of genotypes with expression measurements that differ from *g* by a single nucleotide. Specifically, for each focal genotype *g*, we computed the mean absolute change in regulation strength caused by single-nucleotide mutations as

$$R_{\mathrm{av}}(g) = \frac{1}{n}\sum_{i=1}^{n}|S_R(g) - S_R(g_i)|,$$

where $g_i$ denotes one of the *n* measured single-nucleotide neighbours of *g*, and $S_R(g)$ denotes the normalised regulation strength of genotype *g*. Larger values of $R_{\mathrm{av}}(g)$ therefore indicate greater mutational sensitivity. To convert this metric into a measure of robustness robustness, we report

$$1 - R_{\mathrm{av}}(g),$$

so that higher values correspond to greater robustness and lower values to greater sensitivity. The metric $1 - R_{\mathrm{av}}(g)$ quantifies the expected magnitude of change in regulation strength following a single-nucleotide substitution, and follows the formulation used in a previous study analysing robustness in a tRNA fitness landscape[11]. Robustness estimates are insensitive to incomplete coverage of the theoretical sequence space, and to the resulting variation in neighbourhood size (**Supplementary Fig. S3**).

#### 6.2 Choice of neighbourhood radius

We focused on single-mutant neighbourhoods (Hamming distance = 1) for our primary analysis because double mutations occur exceedingly rarely on short evolutionary timescales in bacteria such as *E. coli*, where the product of effective population size and mutation rate is small ($N\mu < 1$)[12,13]. Single-mutant neighbourhoods are therefore the relevant unit for assessing robustness on the timescale of adaptive evolution in the laboratory [14,15].

### 7. Classification of genotypes

#### 7.1 Peaks and plateaus

A peak is a genotype whose single-nucleotide neighbours all have lower regulation strength than itself. Two peaks connected by an edge of equal regulation strength are individually counted as peaks but together form a "plateau". The genotype with the highest regulation strength in the landscape is the summit or global peak.

#### 7.2 High peaks

High peaks are peaks whose regulation strength exceeds that of the wild-type sequence for the corresponding TF. We also tested alternative definitions — peaks within the top 1% and the top 5% of all regulation strengths — both of which yielded qualitatively identical results.

#### 7.3 Regulation strength classes

For analyses stratified by regulation strength (e.g., of the distribution of mutational effects by regulation-strength class), we assigned genotypes to three classes based on tertiles of the regulation strength distribution within each landscape: low (bottom third), medium (middle third), and high (top third).

### 8. Pairwise genetic distances between high peaks

For each landscape, we computed all pairwise Hamming distances among high-peak genotypes. As a control, we sampled the same number of random genotypes (not restricted to high peaks) from the landscape and computed their pairwise distances. We repeated this random sampling 1,000 times, generating a null distribution against which to compare the observed high-peak distances.

For visualisation (main-text Fig. 3b), we plotted overlapping histograms of the pooled high-peak pairwise distances (red) and the pooled random-sample pairwise distances (grey) for each TF, with vertical lines at the respective mean distances. To identify statistically significant distance differences, we compared the full distribution of high-peak pairwise distances against the pooled distribution of random-sample pairwise distances using a Wilcoxon rank-sum test.

### 9. Simulated adaptive walks

#### 9.1 Model

We simulated adaptive evolution using Kimura's fixation probability[16–18]. For a mutation from genotype $i$ to a neighbouring genotype $j$, we defined the selection coefficient as $s_{ij} = S_R(j) - S_R(i)$,

where $S_R$ is normalised regulation strength. Positive values of $s_{ij}$ correspond to mutations that increase regulation strength, whereas negative values correspond to mutations that decrease regulation strength under the selective regime we consider. We calculated the fixation probability of a mutation as

$$f_{ij} = \frac{1 - e^{-2s}}{1 - e^{-2Ns}}$$

where $f_{ij}$ is the probability that mutation $j$ in the background of genotype $i$ becomes fixed, and $N$ is the effective population size [16–18]. For $s_{ij} = 0$, we used the neutral limit $f_{ij} = 1/N$.

Simulations followed the strong selection, weak mutation (SSWM) regime[14,15], in which populations are monomorphic most of the time. That is, a new allele created by mutation either fixes or becomes extinct before another arises. This regime is appropriate for *E. coli*, where $N \approx 1.8 \times 10^8$ and $\mu \approx 2 \times 10^{-10}$ (yielding $N\mu < 1$)[19]. In the SSWM regime, adaptive evolution can be modelled as an adaptive random walk. Because fixation probabilities are finite for beneficial, neutral, and deleterious mutations, these "Kimura walks" can include steps that reduce regulation strength, especially at smaller population sizes.

#### 9.2 Mutation bias

We incorporated experimentally determined mutation biases for *E. coli* into our evolutionary simulations [20,21]. That is, different types of single-nucleotide substitutions (transitions vs. transversions, and specific base changes) have different probabilities of occurrence, reflecting known biases in bacterial mutation spectra [20,21].

#### 9.3 Simulation procedure

For each landscape, we randomly selected 10,000 distinct non-peak genotypes as starting points for adaptive walks. From each starting genotype, we performed 10,000 independent Kimura walks, each with up to 10 mutation-fixation steps, yielding $10^8$ walks per landscape and population size (**Section 9.4**).

During each walk, starting from a focal genotype $i$, the next genotype with a normalised step probability that combines mutation bias and Kimura's fixation probability. Specifically, for each single-mutant neighbour $j$ of $i$, we computed

$$\pi_{ij} = \frac{\mu_{ij} f_{ij}}{\sum_{k \in \mathcal{N}(i)} \mu_{ik} f_{ik}},$$

where $\mu_{ij}$ is the mutation-bias weight for the substitution from $i$ to $j$ (**Section 9.2**), $f_{ij}$ is the Kimura fixation probability (**Section 9.1**), and $\mathcal{N}(i)$ is the set of single-mutant neighbours of genotype $i$ for which we have measured regulation strength. This set may include beneficial, neutral, and deleterious neighbours, whose fixation probabilities differ according to their selection coefficients and the effective population size $N$[17]. We sampled the next genotype in a single draw from the multinomial distribution defined by $\pi_{ij}$, using the `numpy.random.choice` function in python.

This normalisation preserves the relative probabilities with which alternative mutations become fixed, given their mutation-bias weights and fixation probabilities. It does not preserve absolute mutational waiting times, but our analyses focus on trajectory endpoints, peak accessibility, and regulation-strength gains rather than waiting times. To accelerate computation, we precomputed fixation probabilities for all neighbouring genotype pairs in each landscape.

Because Kimura walks allow neutral and deleterious mutations to fix by genetic drift, they do not necessarily terminate when they reach a local peak. We therefore allowed each walk to proceed for up to 10 mutation-fixation steps. In practice, at $N = 10^8$, most walks stop changing regulation strength substantially within a few steps and reached their last accepted high-regulation state well before this limit (**Supplementary Fig. S6**). The 10-step limit therefore does not bias our accessibility estimates.

#### 9.4 Population sizes

We primarily simulated Kimura walks at $N = 10^8$, but also assessed the sensitivity of our results to decreased population sizes ($N = 10^5$ and $N = 10^2$). Because Kimura fixation probabilities depend on *N*, smaller populations experience stronger genetic drift and have a higher probability of fixing neutral or deleterious mutations. Larger populations, in contrast, are dominated by mutation-fixation events that increase regulation strength.

#### 9.5 Accessibility metrics

We used the following three metrics to quantify peak accessibility.

- **Fraction of walks reaching a high peak**: the proportion of adaptive walks whose endpoint is a high peak (main-text Fig. 3d).
- **Distribution of endpoint regulation strengths**: the cumulative distribution function of regulation strengths at walk endpoints, compared among TFs (main-text Fig. 3c).
- **Number of steps to first high peak**: for walks that do reach a high peak, the number of mutational steps before first arrival to the high peak.

### 10. Randomised null model for accessibility

To test whether observed peak accessibility exceeds what would be expected from genotype-network topology alone, we constructed randomised landscapes by permuting regulation-strength values across genotypes while keeping the genotype graph, including nodes and edges, unchanged. This procedure preserves the mutational neighbourhood structure and the empirical distribution of regulation strengths, but destroys the association between TFBS sequence and regulation strength.

For each TF, we generated 1,000 randomised landscapes in this way. We repeated the adaptive-walk simulations on each randomised landscape using the same starting genotypes, population size, mutation-bias model, and step limit as in the empirical landscape. For each peak accessibility metric, we compared the observed value with the distribution of values obtained from the randomised landscapes. We used one-sided permutation tests to test the null hypothesis that observed accessibility is no greater than expected under random assignments of regulation strength to genotypes. We calculated empirical P-values as the fraction of randomised landscapes with accessibility greater than or equal to the observed accessibility. We summarised results from the randomised-landscape ensemble using the 2.5th and 97.5th percentiles of the corresponding metric

### 11. Distribution of mutational effects (DFE) analysis

For each single-nucleotide mutation in each landscape, we computed the change in regulation strength as

$$\Delta S_R = S_R(j) - S_R(i),$$

where $i$ is the source genotype and $j$ is the single-mutant neighbouring genotype. We characterised the distribution of mutational effects for each TFBS landscape by computing the following quantities:

- **Overall DFE**: density curves of $\Delta S_R$ across all single-nucleotide mutations in each landscape.
- **Class-specific DFE**: separate distributions for mutations originating from genotypes in the low, medium, and high regulation-strength classes, defined by tertiles of $S_R$ within each landscape (**Section 7.3**), and visualised as ridgelines in **Supplementary Fig. S4**.

- **Summary statistics**: mean mutational effect, variance of mutational effects, and proportion of mutations that reduce regulation strength ($\Delta S_R < 0$), computed for each regulation-strength class and TFBS landscape.

We compared distributions of mutational effects between pairs of landscapes using two-sample Kolmogorov–Smirnov tests, applied separately within each regulation-strength class. The null hypothesis was that the mutational effects in each pair of landscapes were drawn from the same distribution.

### 12. Statistical comparisons across TF classes

#### 12.1 Robustness across TF classes

To test whether mutational robustness differs among landscapes, we applied the Kruskal–Wallis rank-sum test[22] to per-genotype robustness values. The null hypothesis was that robustness values are drawn from the same distribution across the five landscapes. We then performed pairwise comparisons between landscapes using Dunn's post-hoc test[23] with Bonferroni corrections for multiple testing.

#### 12.2 Peaks versus non-peaks

We tested whether peak genotypes differ from non-peak genotypes in mutational robustness within each landscape using two-sided Wilcoxon rank-sum tests. The null hypothesis was that robustness values of peak and non-peak genotypes are drawn from the same distribution. We report effect sizes as Cliff's $\delta$, a non-parametric measure of stochastic dominance[24]. Cliff's $\delta$ ranges from $-1$ to +1; negative (positive) values indicate that peak genotypes have lower (higher) robustness than non-peak genotypes.

#### 12.3 High-peak distance comparisons

We assessed differences in pairwise Hamming distances between high peaks and random genotypes (**Section 8**) using Wilcoxon rank-sum tests.

### 13. Position weight matrix (PWM) analysis

To assess the distribution of predicted binding affinities for global TFs across the *E. coli* genome, we extracted manually curated genomic TFBSs from RegulonDB[3] for CRP ($N = 448$ sites), Fis ($N = 327$ sites), and IHF ($N = 152$ sites). For each TF, we constructed a position weight matrix from validated binding sites using the *Biostrings*[25] package in R, with $\log_2$-probability-ratio scoring and uniform background frequencies ($P(A) = P(C) = P(G) = P(T) = 0.25$). We scored each genomic binding site against its cognate PWM by scanning both strands and retaining the highest score. We normalised scores to range from 0 to 1 for comparison with regulation strengths derived from our Sort-Seq experiments.

We used the same PWM construction to identify the eight most informative positions for each TFBS library. Specifically, we calculated information content at each motif position from curated genomic binding sites for the corresponding TF and selected the eight positions with the highest information content, equivalent to the lowest base-frequency entropy. We then randomized these positions in the Sort-Seq libraries (**Section 2.2**).

### 14. Evolvability-enhancing (EE) mutation classification

#### 14.1 Framework

We classified single-nucleotide mutations as evolvability-enhancing (EE), evolvability-neutral (EN), or evolvability-reducing (ED), following the framework of Wagner[26]. For each mutation from a parental genotype $p$ to a single-mutant genotype $m$, we computed the following quantities:

- $\bar{S}_R(n_p)$: the mean regulation strength of the single-mutant neighbourhood of the parental genotype $p$, excluding neighbours that differ from $p$ at the same nucleotide position as the mutation from $p$ to $m$. This exclusion prevents the focal mutation itself, and the two alternative mutations at the same position, from influencing the neighbourhood mean.
- $\bar{S}_R(n_m)$: the analogous mean regulation strength of the single-mutant neighbourhood of the mutant genotype $m$, excluding neighbours that differ from $m$ at the same nucleotide position as the mutation from $p$ to $m$. This also excludes the parental genotype $p$ itself.

We define the direct effect of the mutation on regulation strength [26] as

$$\Delta S_R = S_R(m) - S_R(p)$$

We define an EE mutation as a mutation m that obeys the relationship

$$E = \bar{S}_R(n_m) - \bar{S}_R(n_p) - \max(0, \Delta S_R)$$

This single inequality covers three possible classes of mutations, as follows.

For **beneficial mutations**, where $\Delta S_R > 0$, an EE mutation must satisfy

$$\bar{S}_R(n_m) - \bar{S}_R(n_p) > \Delta S_R$$

Thus, the mutation must increase the mean regulation strength of the mutant neighbourhood by more than expected from its own direct effect on regulation strength. Under additivity, the expected relationship is $\bar{S}_R(n_m) = \bar{S}_R(n_p) + \Delta S_R$. The EE criterion therefore identifies mutations that, on average, interact positively with subsequent mutations in their local neighbourhood[26].

For **deleterious mutations**, where $\Delta S_R < 0$, an EE mutation must satisfy

$$\bar{S}_R(n_m) - \bar{S}_R(n_p) > 0.$$

Thus, even though the focal mutation itself reduces regulation strength, the single-mutant neighbourhood of the mutant genotype must have higher mean regulation strength than the corresponding neighbourhood of the parental genotype. This stricter criterion avoids classifying deleterious mutations as EE merely because subsequent mutations partially compensate for their deleterious effect. Instead, deleterious EE mutations must shift the local neighbourhood toward higher regulation strength[26].

Strictly neutral EE mutations, where $\Delta S_R = 0$, were not enumerated as a separate class because reliable identification of neutrality from Sort-Seq data is limited by measurement noise. Mutations with effects sufficiently close to zero were therefore assigned to the beneficial or deleterious class according to the sign of their estimated $\Delta S_R$.

**14.2 Statistical test**

For both beneficial and deleterious classes we used a two-sided one-sample *t*-test of the unified null hypothesis that

$$E = \bar{S}_R(n_m) - \bar{S}_R(n_p) - \max(0, \Delta S_R)$$

differs significantly from zero, where $\Delta S_R = S_R(m) - S_R(p)$. We used a two-sided one-sample (t)-test with the null hypothesis that $E = 0$. The test statistic was

$$t = \frac{\bar{S}_R(n_m) - \bar{S}_R(n_p) - \max(0, \Delta S_R)}{\sigma_E}$$

where $\sigma_E$ is the standard error of the neighbourhood effect. We estimated $\sigma_E$ by propagating per-genotype Sort-Seq replicate variance through the difference between neighbourhood means as

$$\sigma_E \approx \sqrt{\frac{1}{n_m^2}\sum_{j=1}^{n_m} \operatorname{var}\left(S_R(m_j)\right) + \frac{1}{n_p^2}\sum_{j=1}^{n_p} \operatorname{var}\left(S_R(p_j)\right)},$$

where $m_j$ and $p_j$ denote valid neighbours of the mutant and parental (pre-mutation) genotypes, respectively, and $n_m$ and $n_p$ are the corresponding neighbourhood sizes. We used $\min(n_m, n_p) - 1$ degrees of freedom, which makes the test conservative when neighbourhood sizes differ.

This formulation differs from that of Wagner[26], who estimated the variance of the neighbourhood effect from the empirical variance among fitness values within each neighbourhood. We instead used per-genotype measurement variance from Sort-Seq replicates (**Section 14.3**), because this directly captures the measurement noise relevant to classifying individual mutations in our regulation-strength landscapes. In contrast, variance among neighbours combines measurement noise with biological differences in regulation strength among neighbouring genotypes, which is the signal the EE classification is designed to detect. Validation against the RNA landscape [11] analysed by Wagner [26] confirmed that the two formulations produce essentially identical classifications when measurement variance and between-neighbour variance have similar magnitude (**Section 14.7**). The relevant degrees of freedom for the test we use equal min($n_p$, $n_m$) − 1, which renders the test conservative when neighbourhood sizes differ.

#### 14.3 Measurement variance

We estimated per-genotype measurement variance from Sort-Seq replicate data. For each genotype $i$, we computed the variance of the normalised regulation strength score as

$$\mathrm{var}_i = \left(\frac{S_R(i),\mathrm{CV}i}{\sqrt{n\mathrm{reps}}}\right)^2,$$

where $S_R(i)$ is the mean normalised regulation strength of genotype $i$, $\mathrm{CV}i$ *is* the genotype's coefficient of variation in regulation strength across biological replicates, and $n\mathrm{reps}$ is the number of biological replicates ($n\mathrm{reps} = 3$ for TetR, CRP, Fis, and IHF, and $n\mathrm{reps} = 4$ for LasR). This variance estimates the uncertainty of the mean regulation-strength value assigned to each genotype, and was used in the replicate-based EE classification test described in **section 14.1**.

#### 14.4 Multiple testing correction

We corrected all *P*-values using the Benjamini–Hochberg procedure[27] at a false discovery rate (FDR) of 1%, applied separately to beneficial and deleterious mutations.

#### 14.5 Randomised null model for EE enrichment

To assess whether observed EE fractions exceed chance expectation, we generated 1,000 randomised landscapes for each TF by permuting regulation-strength values across genotypes while preserving genotype-network topology, as described in **Section 10**. We applied the EE classification procedure (**Sections 14.1–14.4**) identically to each randomised landscape and compared the observed EE fraction with the distribution of EE fractions obtained from the randomised landscapes. We computed fold enrichment as the ratio between the EE fraction in the empirical landscape and the mean EE fraction across randomised landscapes. We used one-sided permutation tests with the null hypothesis that the observed EE fraction is no greater than expected under random assignment of regulation strength to genotypes. We calculated empirical P-values as the fraction of randomised landscapes with a fraction of EE mutations greater than or equal to the observed fraction.

#### 14.6 Coverage

We performed EE classification exhaustively on every single-nucleotide mutation in each landscape's giant component (TetR: $n$ = 14,458 source genotypes after exclusion of genotypes lacking valid neighbourhood variance estimates; LasR: 43,076; CRP: 31,811; Fis: 43,210; IHF: 41,296). We applied no subsampling of mutations for different landscapes. Earlier exploratory analyses on a 1,000-genotype random subsample (used during code development) yielded EE fractions within one percentage point of the full-landscape values, indicating that the full-landscape numbers are not driven by sampling artefacts.

#### 14.7 Validation

We validated our implementation with the procedure used by Wagner[26] to analyze the tRNA fitness landscape of Domingo et al.[11,] and the protein toxin–antitoxin landscape of Lite et al.[28]. Our implementation yielded an overall EE fraction of 12.71% (matching the value of 12.71%, reported by Wagner[26]), Benjamini–Hochberg[27] $P$-value thresholds of 0.00645/0.00635 (reported value: 0.00640), and full classification agreement of 99.9%, confirming faithful reproduction of the published analysis.

#### 14.8 Consequences for adaptive walks

To evaluate whether EE mutations influence adaptive-walk outcomes, we partitioned the Kimura-walk simulations (Section 9) into two classes:

- **EE-mutation containing walks**, which included at least one mutational step classified as EE ($n_{\mathrm{EE}} \geq 1$) in the walk-level summary.
- **Non-EE-mutation containing walks**, which contained only EN or ED steps.

For each walk, we calculated the regulation-strength gain as $G_R = S_R^{\mathrm{endpoint}} - S_R^{\mathrm{start}}$, where $S_R^{\mathrm{start}}$ and $S_R^{\mathrm{endpoint}}$ are the normalised regulation strengths of the starting and endpoint genotypes, respectively. We compared $\Delta G_R$between EE-mutation containing and non-EE-mutation containing walks using two-sided Mann–Whitney U tests. The null hypothesis was that regulation-strength gains are drawn from the same distribution for the two walk classes. We performed this comparison at the primary population size N = $10^8$ (main-text Fig. 4d) and repeated it at N = $10^5$ and N = $10^2$ to test whether the within-walk benefit of EE mutations persists under stronger genetic drift (Supplementary Fig. S12). We also recorded whether each walk terminated at a local peak below the high-peak threshold (Section 7.2), recorded total walk length, and assessed the association between EE-step count and walk length using Spearman's rank correlation.

#### 14.9 Fraction of beneficial neighbours and adjusted neighbourhood mean

To examine how an EE mutation affects the local mutational neighbourhood of the mutant genotype, we computed three quantities for each EE mutation, following ref.[26]:

- **Mean neighbour regulation strength:** the mean regulation strength of the single-mutant neighbourhood of the parental (pre-mutation) genotype, $\bar{S}_R(n_p)$, and of the EE mutant genotype, $\bar{S}_R(n_m)$. These paired distributions are shown in Supplementary Fig. S7.

- **Adjusted mean neighbour regulation strength:** $\bar{S}_R(n_p) - S_R(p)$ and $\bar{S}_R(n_m) - S_R(m)$, where $p$ is the parental genotype and $m$ is the EE mutant genotype. This adjustment subtracts each focal genotype's own regulation strength, allowing us to compare how the neighbourhood mean differs from the focal genotype before and after the EE mutation. These distributions and paired comparisons are shown in **Supplementary Fig. S8**.
- **Fraction of beneficial neighbours:** the proportion of single-mutant neighbours of the parental genotype and of the EE mutant genotype whose regulation strength exceeds that of the parental genotype. These distributions are shown in main-text **Fig. 4c**.

For each TF, we used paired Wilcoxon signed-rank tests to test the null hypothesis that parental genotypes and their corresponding EE mutants have the same median fraction of beneficial neighbours. This analysis tests whether EE mutations increase the number of beneficial one-step mutations in a genotype's neighbourhood. Because EE mutations are defined by their effect on neighbourhood mean regulation strength relative to an additive expectation, an increased fraction of beneficial neighbours provides an intuitive summary of how EE mutations can alter subsequent adaptive-walk trajectories.

#### 14.10 Within-landscape coupling between robustness and EE mutation prevalence

For each landscape, we computed two genotype-level quantities: mutational robustness (Section 6.1) and the fraction of EE mutations of each focal genotype. We calculated this EE fraction separately for three mutation sets: all mutations, beneficial mutations only, and deleterious mutations only (Supplementary Fig. S11). We then tested whether genotypes with higher robustness also tended to have higher or lower fractions of EE mutations using Spearman's rank correlation coefficient. For each landscape and mutation set, the null hypothesis was that genotype-level robustness and genotype-level EE fraction are not monotonically associated.

After excluding genotypes lacking valid EE classifications, within-landscape sample sizes in these tests were n = 14,458 for TetR, 43,076 for LasR, 31,811 for CRP, 43,210 for Fis, and 41,296 for IHF. We calculated Spearman's $\rho$ and associated P-values using the `cor.test` function in R (version 4.3.0).

#### 14.11 Cross-landscape relationship between robustness and EE mutation prevalence

We also tested whether EE mutation prevalence covaries with mean mutational robustness across the five TFBS landscapes. To this end, we computed for each landscape two landscape-level quantities: the mean genotype robustness for all genotypes in the landscape (**Section 6.1**), and the overall fraction of beneficial mutations classified as EE (**Section 14.1**). We then quantified the association between the resulting five pairs of values using Spearman's rank correlation coefficient, and tested the null hypothesis that mean landscape robustness and beneficial-EE prevalence are not monotonically associated across landscapes (Spearman's $\rho = -0.10$, P = 0.95; **Supplementary Fig. S9**).

### 15. Software and computational environment

We performed all genotype network construction and adaptive walk simulations using in-house Python (version 3.13) scripts with the *igraph*[9] and *NumPy* packages[29]. We performed robustness calculations, EE classification, statistical analyses, and visualisations in R (version 4.3 or higher) using the following packages: *igraph*, *tidyverse* (including *dplyr*, *ggplot2*, *tidyr*, *stringr*, *readr*, *purrr*), *patchwork*, *ggrepel*, *ggridges*, *ggpubr*, *scales*, *viridis*, *Biostrings*, *ggraph*, *tidygraph*, and *Cairo*. All code is available in the GitHub repository accompanying this paper.

### Supplementary Figures

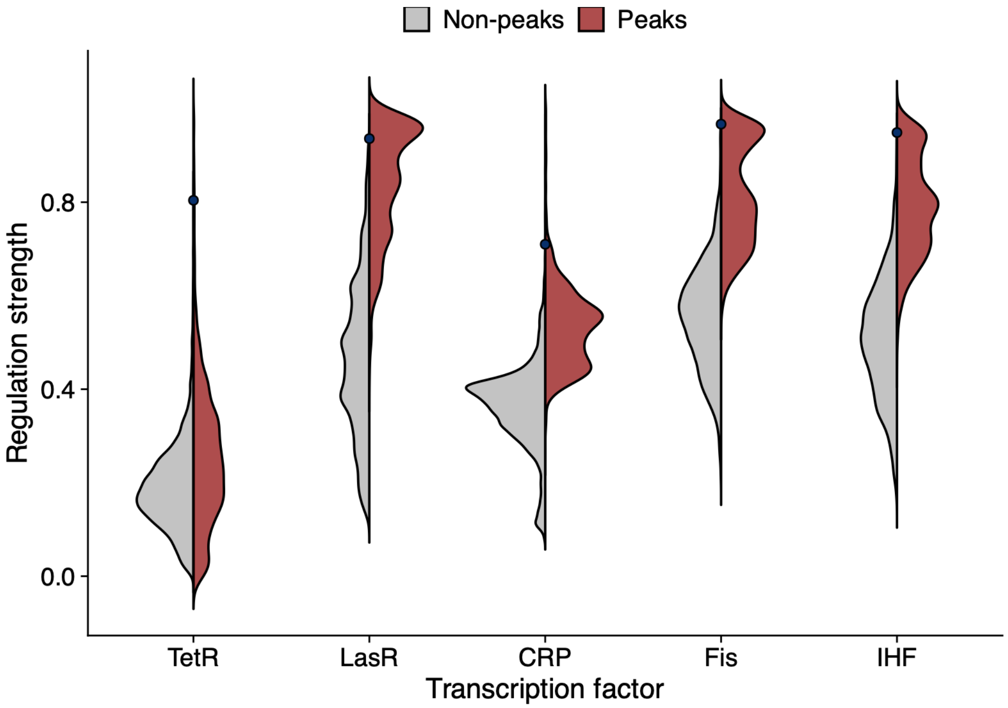


**Figure S1. Distribution of regulation strengths across landscapes.** Violin plots depict the range of binding strengths for TFBS variants of different transcription factors (TetR, LasR, CRP, Fis, and IHF). Red regions indicate peak variants, while gray regions represent non-peak variants. The blue circles indicate the wild-type regulation strength as a baseline for categorizing high peaks (above the WT)

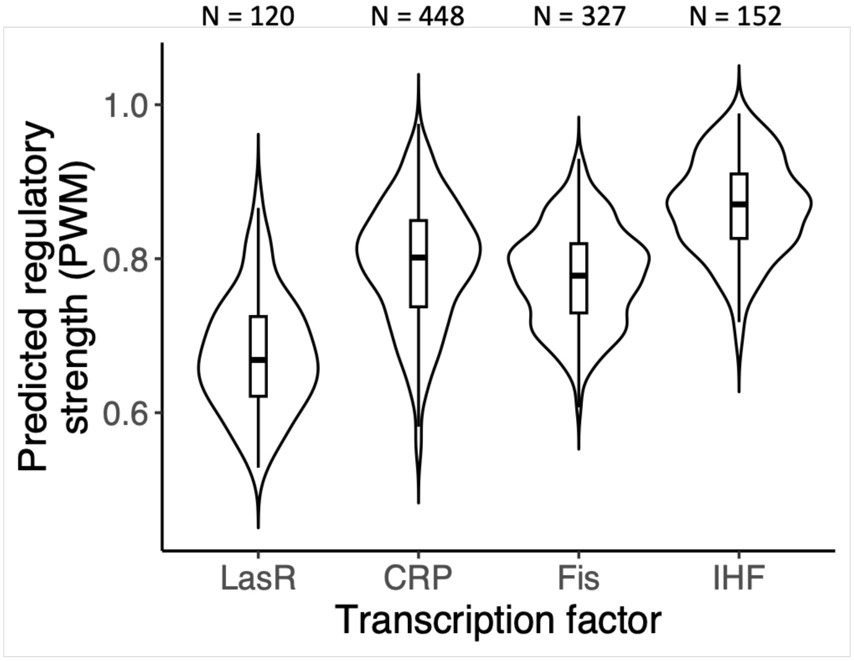


**Figure S2. Distribution of predicted regulation strengths for genomic TFBSs.** Violin plots show the distribution of position weight matrix (PWM) scores for LasR (orange, $N$ = 120 sequences), CRP (green, $N$ = 448 sequences), Fis (blue, $N$ = 327 sequences), and IHF (purple, $N$ = 152 sequences) binding sites (**Methods**). For CRP, Fis, and IHF, we extracted manually curated *E. coli* genomic TFBSs from the RegulonDB database[3]. For LasR, we used experimentally validated *P. aeruginosa* binding sites from ref.[30]. For each transcription factor, we constructed a PWM from validated binding sites and used it to compute predictive PWM scores. A PWM represents the binding preferences of a transcription factor by capturing the probability of each nucleotide at each position within a motif, derived from experimentally validated binding sites[31,32]. These probabilities are normalised and converted into log-odds scores, highlighting sequence positions enriched for specific nucleotides relative to the background nucleotide frequencies. The distributions reflect the variability in predicted regulation strengths across TFBSs for each transcription factor.

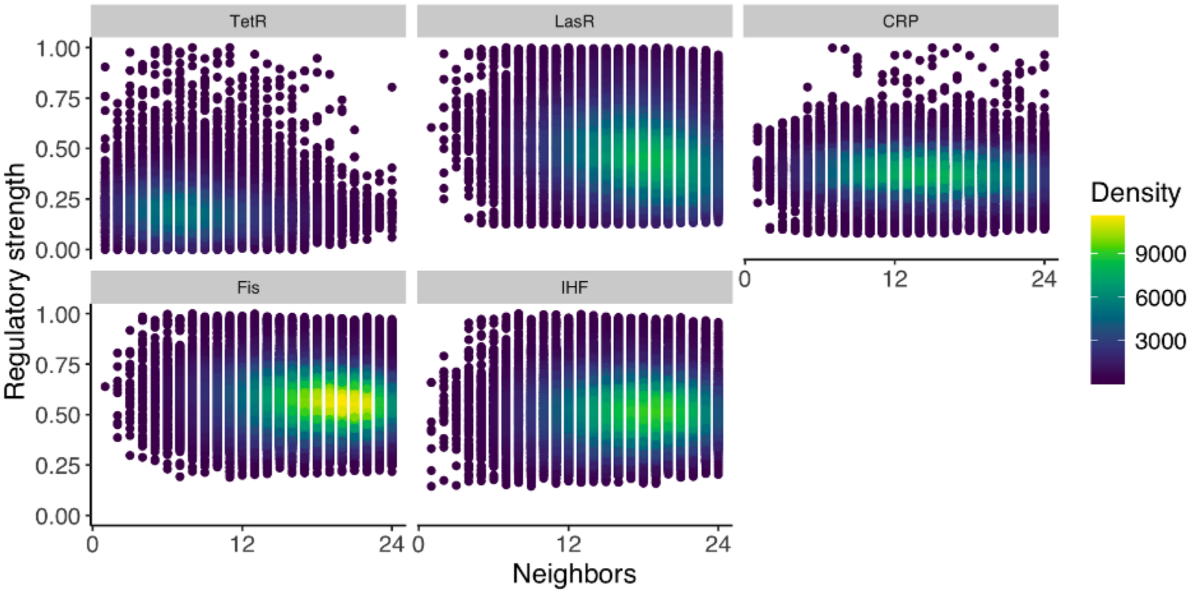


**Figure S3. Relationship between the number of neighbours and regulation strength across TFBS landscapes.** Scatter density plots show the number of single-nucleotide neighbors (x-axis) versus normalized regulation strength (y-axis) for TetR (a), LasR (b), CRP (c), Fis (d), and IHF (e) TFBS landscapes. Each circle represents a genotype, and color intensity reflects local density of data. The number of neighbors corresponds to the number single-nucleotide variants with experimental Sort-Seq regulation strength measurements for each genotype, with a maximum of 24 (three alternatives at each of eight randomised positions). Genotypes span the full range of regulation strengths at nearly all neighborhood sizes, indicating that incomplete neighborhood coverage does not introduce systematic biases in the estimation of robustness or other network-derived metrics.

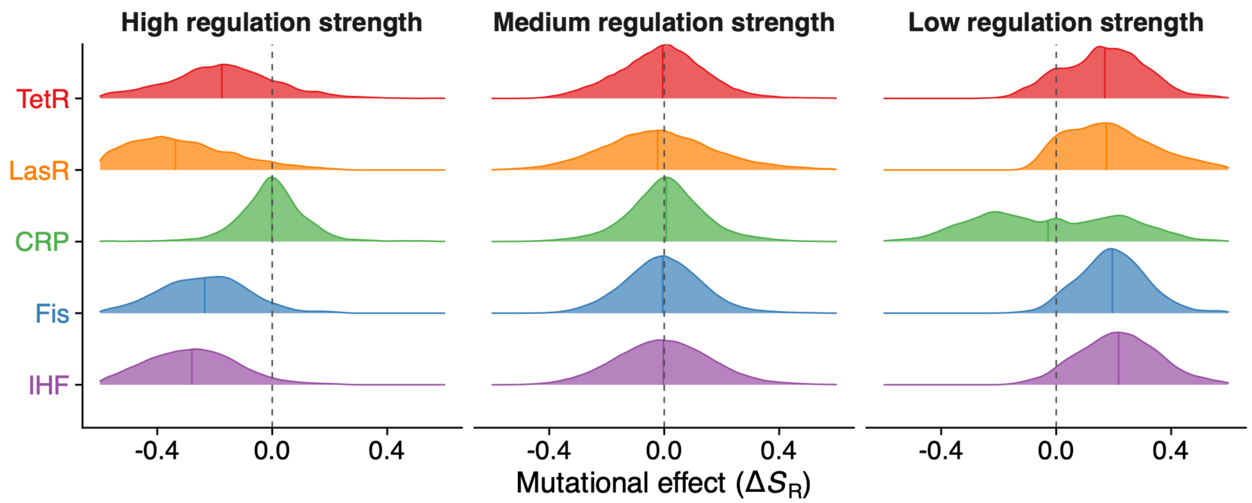


**Supplementary Figure S4. Distribution of mutational effects (DFE) by regulation-strength class.** Ridge plots showing the distributions of single-nucleotide mutational effects on regulation strength, defined as $\Delta S_R = S_R(j) - S_R(i)$, where $i$ is the focal genotype and $j$ is its single-mutant neighbour. Distributions are shown for the five TFBS landscapes (rows; colour-coded as elsewhere) across three regulation-strength classes (columns), defined as tertiles of the within-landscape regulation-strength distribution (**Supplementary Methods Section 7.3**): high (top tertile), medium (middle tertile), and low (bottom tertile). The dashed vertical line at 0 indicates no mutational effect, and the coloured vertical line within each ridge plot marks the mean of the distribution. Across landscapes, mutational effects are biased toward decreases in regulation strength in the high-strength class and toward increases in the low-strength class, with the strongest asymmetry in the TetR landscape. Sample sizes (number of mutations per TFBS landscape × class combination) range from $3.0 \times 10^4$ for TetR in the low-strength class to $1.3 \times 10^5$ for Fis in the medium-strength class.

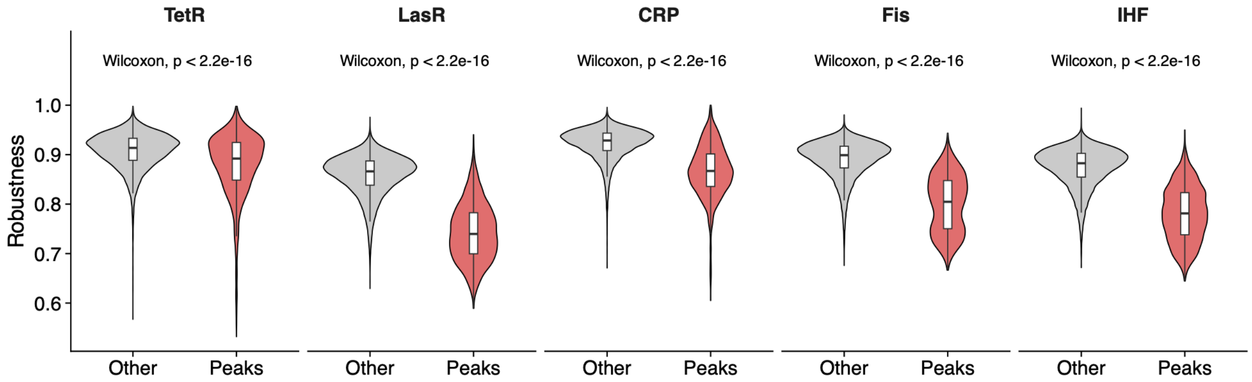


**Supplementary Figure S5. Peak genotypes are mutationally less robust than non-peak genotypes in every landscape.** Distribution of robustness ($R_{avg}$) for peak (red) versus non-peak (grey) genotypes in each landscape. Box plots inside the violins show the interquartile range (IQR) with the median marked by a horizontal line and whiskers extending to 1.5× IQR. P-values are based on two-sided Wilcoxon rank-sum tests. Cliff's δ effect sizes are −0.45 (TetR), −0.88 (LasR), −0.65 (CRP), −0.81 (Fis), and −0.83 (IHF), all consistent with peaks being less robust than non-peak genotypes. Sample sizes: n_peaks = 2,068 (TetR), 2,580 (LasR), 2,132 (CRP), 2,327 (Fis), 2,450 (IHF); n_non-peaks = 13,388, 40,506, 29,801, 40,894, 38,867 respectively.

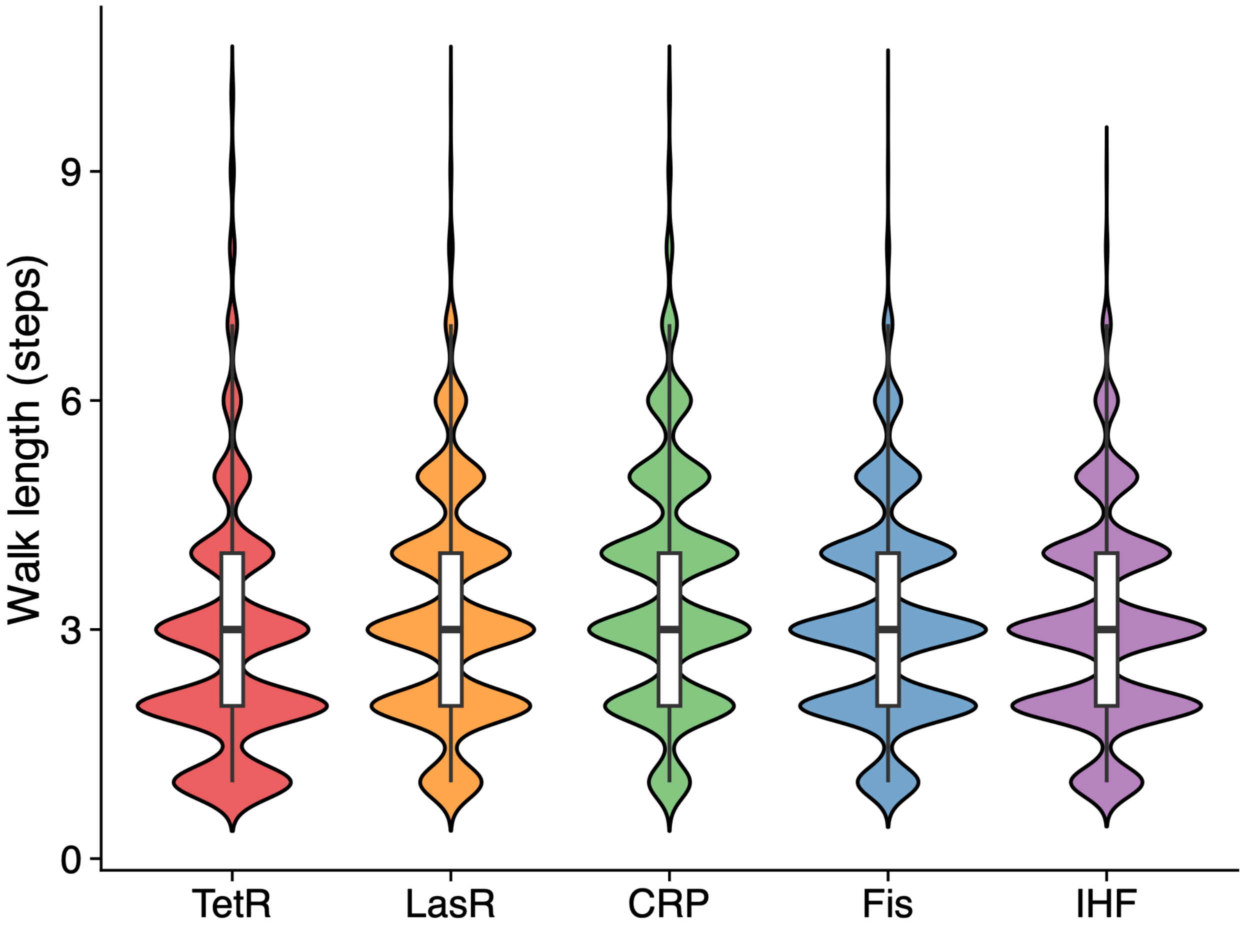


**Supplementary Figure S6. Adaptive walks settle quickly: most reach a local peak within 3–5 mutational steps.** Distribution of walk lengths — defined as the mutation-fixation step at which the walk last accepted a mutation — across all $10^8$ adaptive walks per landscape at a population size N = $10^8$ (**Methods, Supplementary Methods Section 9.3**). Violins show kernel densities; box plots inside show the IQR with the median marked by a horizontal line, and whiskers extending to 1.5× IQR. The median walk length is 3 steps for all five landscapes; 95% of walks complete within 7 steps. The 10-step ceiling imposed during simulation is therefore well above the observed walk lengths, so it does not bias accessibility estimates.

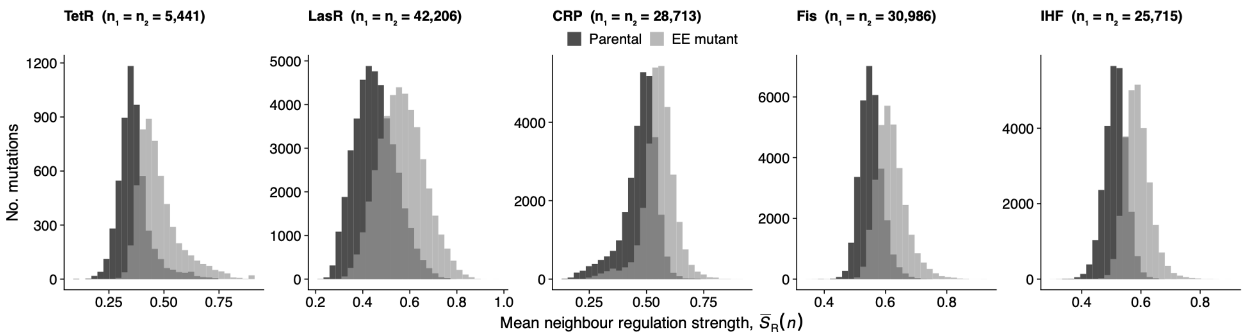


**Supplementary Figure S7. EE mutants have higher mean neighbour regulation strength than their parental genotypes.** Histograms of the mean regulation strength of single-mutant neighbours, $\bar{S}_R(n)$, for parental genotypes (dark grey) and their corresponding EE mutant genotypes (light grey), shown separately for each TFBS landscape. Vertical lines mark the means of each distribution. Sample sizes are matched between groups by pairing each EE mutant with its parental genotype: $n_1 = n_2 = 5{,}441$ for TetR, 42,206 for LasR, 28,713 for CRP, 30,986 for Fis, and 25,715 for IHF. Across all five landscapes, EE mutants have systematically higher mean neighbour regulation strength than their parental genotypes.

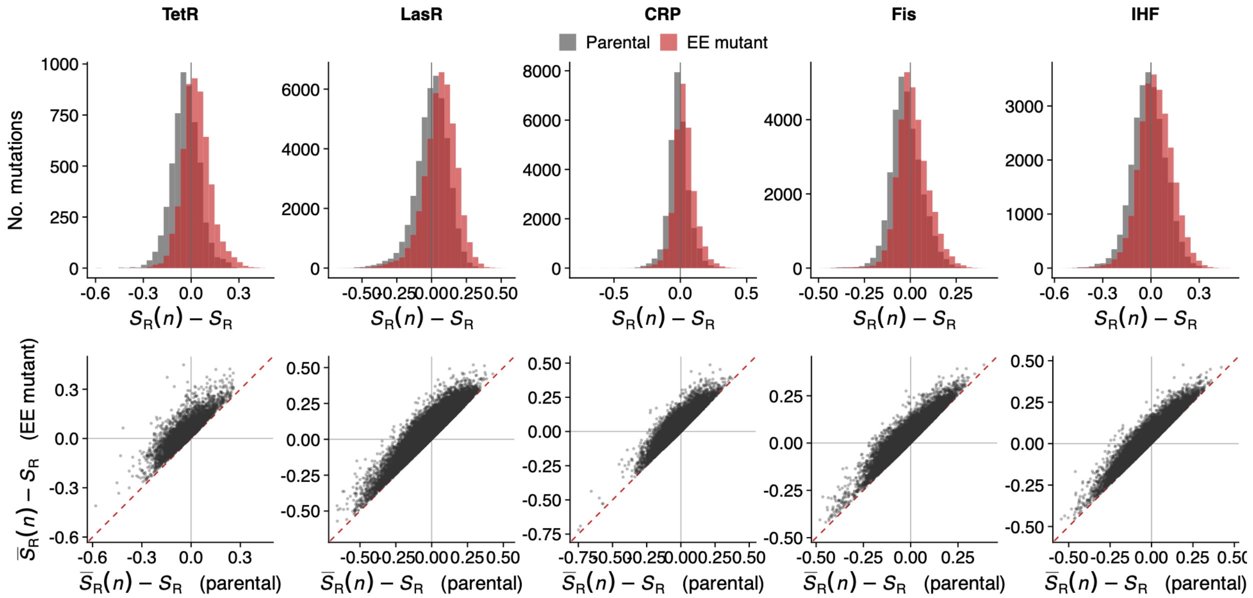


**Supplementary Figure S8. EE mutants increase adjusted mean neighbour regulation strength.** Top row: histograms of adjusted mean neighbour regulation strength for parental genotypes (grey) and their corresponding EE mutant genotypes (red), shown separately for each TFBS landscape. Adjusted mean neighbour regulation strength is defined as $\bar{S}_R(n_p) - S_R(p)$ for the parental genotype and $\bar{S}_R(n_m) - S_R(m)$for the EE mutant genotype, where $\bar{S}_R(n)$ is the mean regulation strength of the single-mutant neighbourhood and $S_R$ is the regulation strength of the focal genotype. Bottom row: paired scatter plots comparing the adjusted mean neighbour regulation strength of each parental genotype (horizontal axis) with that of its corresponding EE mutant genotype (vertical axis). The dashed red line indicates the identity line $y = x$. Each point represents one EE mutation paired with its parental genotype; sample sizes are identical to those in Supplementary Fig. S7. Points above the identity line indicate that the EE mutant has a higher neighbourhood mean relative to its own regulation strength than the parental genotype does.

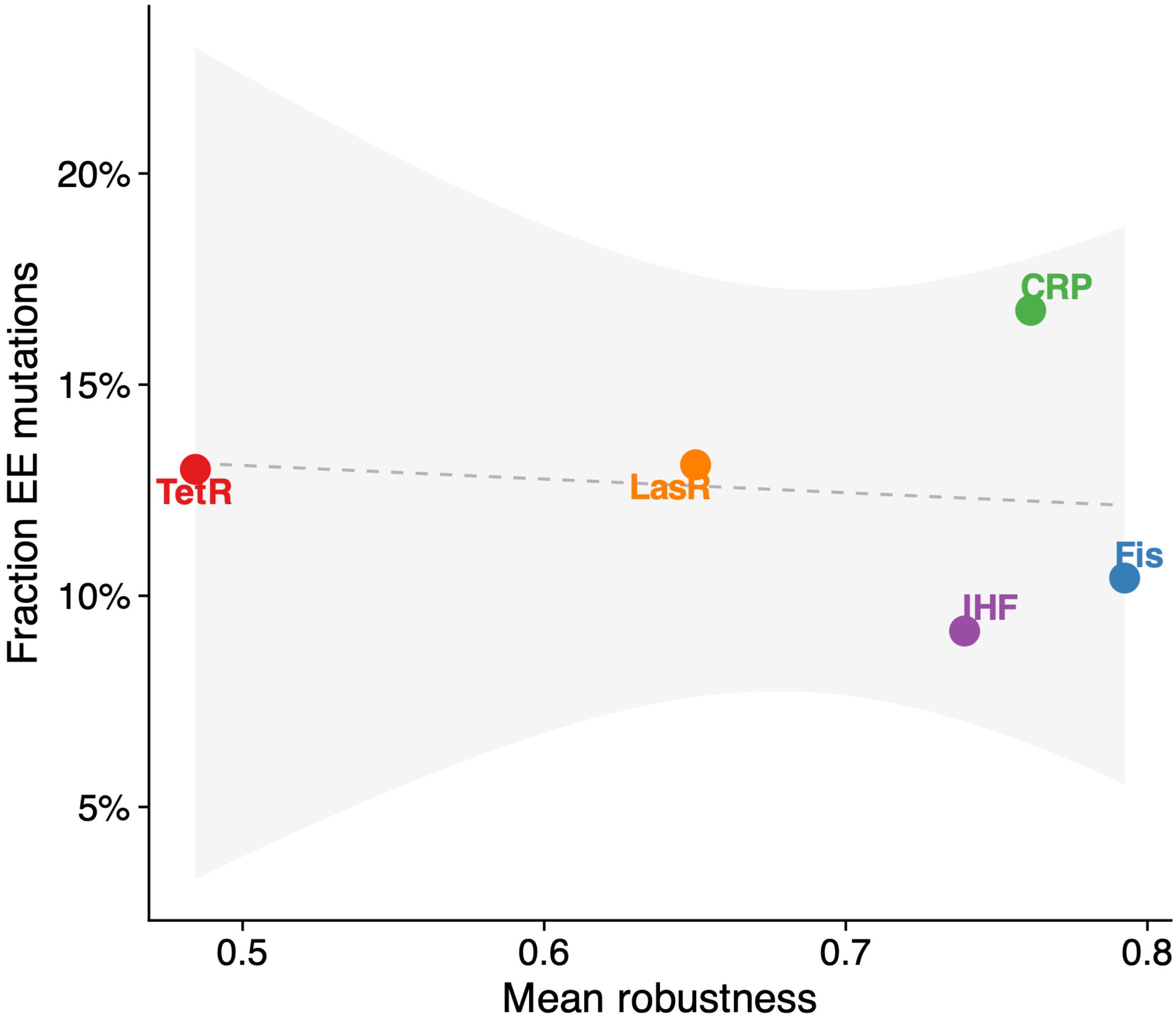


**Supplementary Figure S9. Cross-landscape relationship between mean robustness and EE prevalence.** Mean robustness versus the overall fraction of beneficial mutations classified as EE, with each landscape shown as one labelled point and coloured according to the main manuscript's colour scheme. The dashed line is the best-fit linear regression line; the 95% confidence interval is shaded in grey. With $n = 5$ landscapes, the test does not have power to detect any but the strongest monotonic relationships; the observed correlation is Spearman $\rho = -0.10$ ($P = 0.95$) and is reported without interpretation in the main text (**Supplementary Methods Section 14.11**).

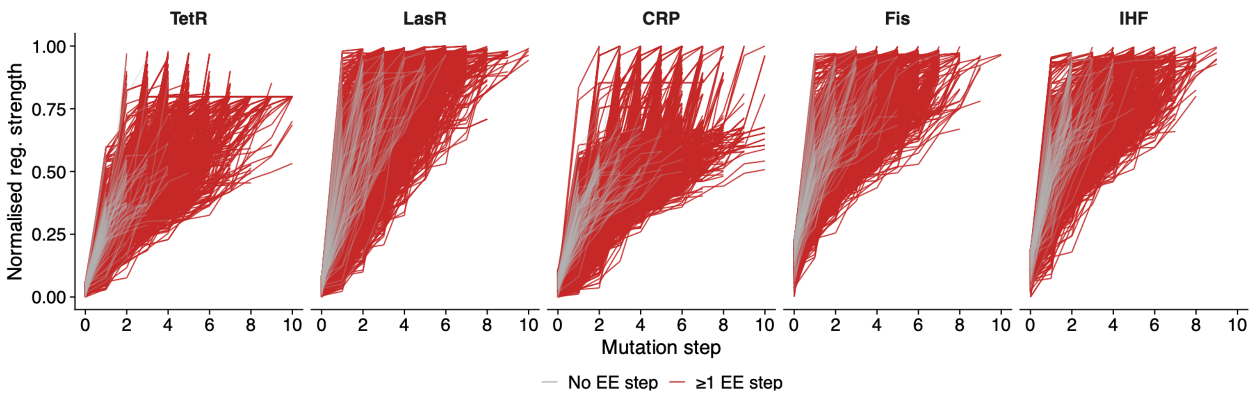


**Supplementary Figure S10. Adaptive walks containing EE mutation-fixation steps reach higher regulation strength across all five landscapes.** Per-walk trajectories of normalised regulation strength as a function of mutational step, for each landscape, at N = $10^8$. Each line is a single adaptive walk, coloured red when it contains at least one EE step and grey when it contains none. All walks containing at least one EE step are shown; to avoid overplotting the far more numerous EE-free walks, a random subsample of up to 100 EE-free walks per landscape is displayed. Walks are short under directional selection — the median walk length is three mutational steps and essentially no walk reaches ten steps (full walk-length distribution in **Supplementary Fig. S6**) — so most trajectories terminate within a few steps. E EE-mutation containing walks (red) systematically reach higher endpoint regulation strengths than walks composed of EN/ED steps only (grey), consistent with the quantitative comparison in main-text **Fig. 4d**.

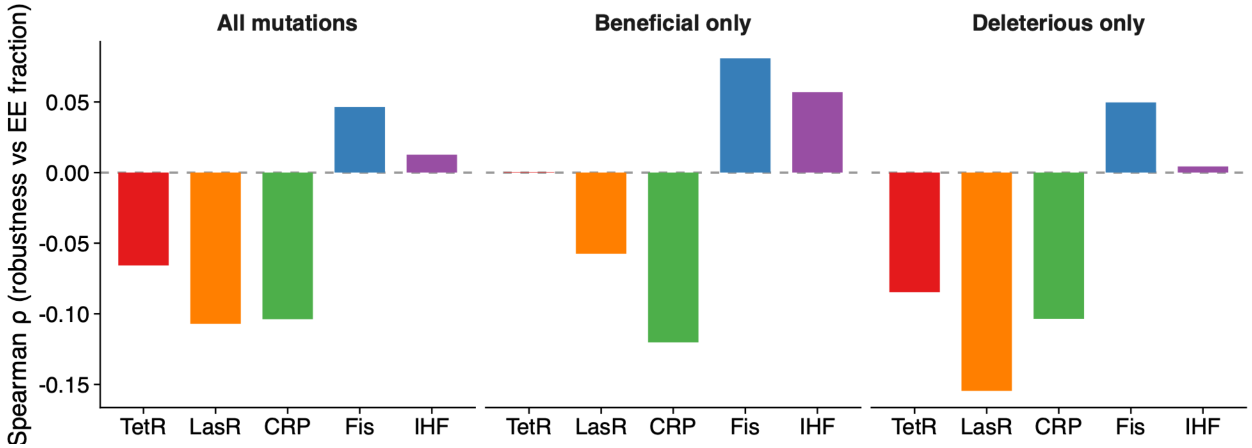


**Supplementary Figure S11. Within-landscape coupling between robustness and EE mutation prevalence is weak and inconsistent in sign.** Per-genotype Spearman rank correlation coefficient $\rho$ between robustness and the fraction of mutations classified as EE, computed separately for all mutations (left), beneficial mutations only (middle), and deleterious mutations only (right). Bars are coloured by TF. Sample sizes per landscape are (n = 14,458) for TetR, 43,076 for LasR, 31,811 for CRP, 43,210 for Fis, and 41,296 for IHF. For all mutations, correlations are ($\rho$ = -0.066) for TetR, (-0.107) for LasR, (-0.104) for CRP, (+0.046) for Fis, and (+0.013) for IHF. Across mutation classes, the sign of the correlation differs among landscapes, and effect sizes are uniformly small $|\rho| < 0.16$

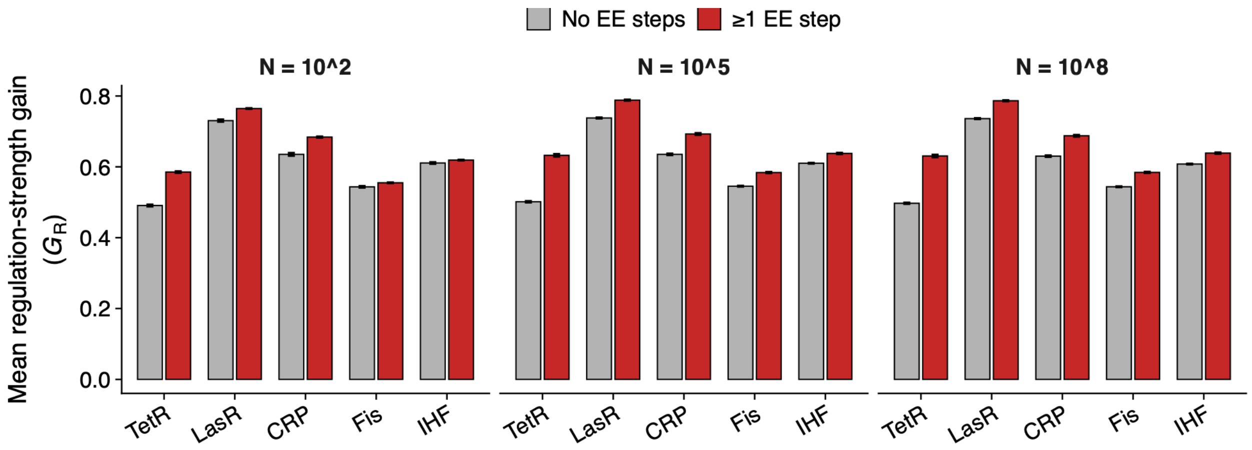


**Supplementary Figure S12. Walks with EE mutations show larger regulation-strength gains at different population sizes.** Mean regulation-strength gain ($G_R$), defined as $S_R^{\text{endpoint}} - S_R^{\text{start}}$, for adaptive walks containing no EE steps (grey) versus walks containing at least one EE step (red). Results are shown for three effective population sizes spanning six orders of magnitude: N = $10^2$, $10^5$, and $10^8$ (**Supplementary Methods Section 9.4**). Error bars indicate one standard error of the mean; in many cases, error bars are smaller than the bars themselves and therefore not visible. For each TF and population size, data are based on $10^8$ walks, corresponding to 10,000 starting genotypes and 10,000 walks per starting genotype, partitioned by EE status as in main-text Fig. 4d. The advantage of EE-mutation containing walks is consistent across all three population sizes, indicating that the within-walk benefit of EE mutations is not restricted to large populations where genetic drift is weak.
